# Auditory cortex conveys non-topographic sound localization signals to visual cortex

**DOI:** 10.1101/2023.05.28.542580

**Authors:** Camille Mazo, Margarida Baeta, Leopoldo Petreanu

## Abstract

Perception requires binding spatiotemporally congruent multimodal sensory stimuli. The auditory cortex (AC) sends projections to the primary visual cortex (V1), which could provide signals for binding spatially corresponding audio-visual stimuli. However, it is unknown whether AC inputs in V1 encode sound location. We used dual-color two-photon axonal calcium imaging and an array of speakers to measure the auditory spatial information that AC transmits to V1. We found that AC relays information about the location of ipsilateral and contralateral sound sources to V1. Sound location could be accurately decoded by sampling AC axons in V1, providing a substrate for making location-specific audiovisual associations. However, AC inputs were not retinotopically arranged in V1, and audio-visual modulations of V1 neurons did not depend on the spatial congruency of the sound and light stimuli. The distributed, non-topographic sound localization signals provided by AC might allow the association of specific audiovisual spatial patterns in V1 neurons.

## INTRODUCTION

Animals probe their environment through multiple specialized sensory organs and integrate them into a coherent percept to guide behavior ^1^. Multimodal sensory stimuli from a common source are congruent in space and time and are bound together into an unified cross modal object ^2^. Sounds can alter our visual perception and vice versa, as experienced in cross-modal illusions such as the ventriloquism effect ^3^, the McGurk effect ^4^ and the double flash illusion ^5^. However, the neuronal mechanisms by which spatially congruent audiovisual stimuli are bound together remain poorly understood.

Visual space is directly mapped at the surface of the retina and information about the spatial location of sound sources, while not directly available in the sensory organs, can be computed using auditory cues ^6^. It is well stablished that spatially congruent auditory and visual stimuli are integrated in the optic tectum of birds and superior colliculus (SC) of mammals^1^. These structures contain spatial auditory and visual maps, and bimodal neurons whose spatial receptive fields (RFs) are in register (barn owl ^7^, ferret ^8^, cat ^6^, guinea pig ^9^, and mouse ^10^). The neural register of auditory and visual representations in these structures provides a substrate for binding spatially congruent auditory and visual stimuli. Consistent with this, auditory and visual stimuli originating from the same location results in larger neural responses in the SC than when they are distant from each other ^11, 12^.

In addition to the SC, it is becoming increasingly evident that sensory areas of the neocortex, even the primary ones, also participate in multimodal sensory integration ^13^. Across species, visual responses in primary visual cortex (V1) are modulated by sounds ^14–23^. Direct projections from auditory cortex (AC) to V1 are also conserved across species ^23–28^ and are thought to mediate at least some of the audiovisual interactions observed in V1 neurons ^15, 17, 21, 29^ (but see ^30^). AC neurons show tuning to sound location ^31–39^. Thus, by sampling AC afferents, neurons in V1, like those of the SC, could potentially have access to both auditory and visual spatial information. While AC does not harbor a topographic map of space ^32, 40^, through selective innervation, spatial information in AC inputs could become aligned with the retinotopic map in V1, as feedback projections from higher visual areas do ^41^. Such an organization could constitute a neural substate for a cortical representation of spatially congruent audiovisual objects.

While neurons projecting to V1 are known to selectively encode sound features that are less represented within AC ^17^, it remains unknown whether they relay information about the location of sound sources, as required for V1 neurons to bind spatially congruent bimodal stimuli by sampling these inputs.

In this study, we measured the auditory spatial information in AC afferents in layer 1 of mouse V1 and compared it to the RF of their postsynaptic neurons. We found that many AC to V1 (AC➔V1) inputs in layer 1 have spatially restricted RFs, encoding spatial information that V1 neurons can potentially use to accurately locate sound sources. However, the auditory RFs in AC inputs in V1 span ipsi- and contralateral hemifields and bear no relation with the visual RFs of their postsynaptic V1 neurons. Consistent with this, we observed that visual responses in V1 neurons are equally enhanced by sounds, regardless of the degree of spatial congruence between the auditory and visual stimuli.

## RESULTS

### AC axons convey auditory spatial information to V1

To gain spatial control over auditory and visual stimuli, we designed an array of loudspeakers and light-emitting diodes (LED) spaced over a spherical section. The array consisted of 39 positions, distributed in 13 columns (10° steps) along the azimuth and 3 rows (20° steps) in elevation (Figure 1a). An LED and a loudspeaker were mounted at each position. We also designed custom electronics boards and software to route signals to specific individual LEDs and speakers in the array (Methods). Mice were head-fixed in the center such that their head was equidistant to all the positions in the array. Mice faced the 3rd column and 2nd row, allowing stimuli to be presented over azimuthal positions spanning from 20° at the hemifield ipsilateral to the recorded hemisphere (left, negative azimuth values) to 100° of the contralateral one, and 20° up and down in elevation relative to the head position (Figure 1b). Thus, the array allowed presenting visual and auditory stimuli in locations spanning most of the azimuthal extent of the visual hemifield represented in V1. We presented interleaved auditory white noise bursts (2-20 kHz) and flashing white LED light (Figure 1c).

**Figure 1.**
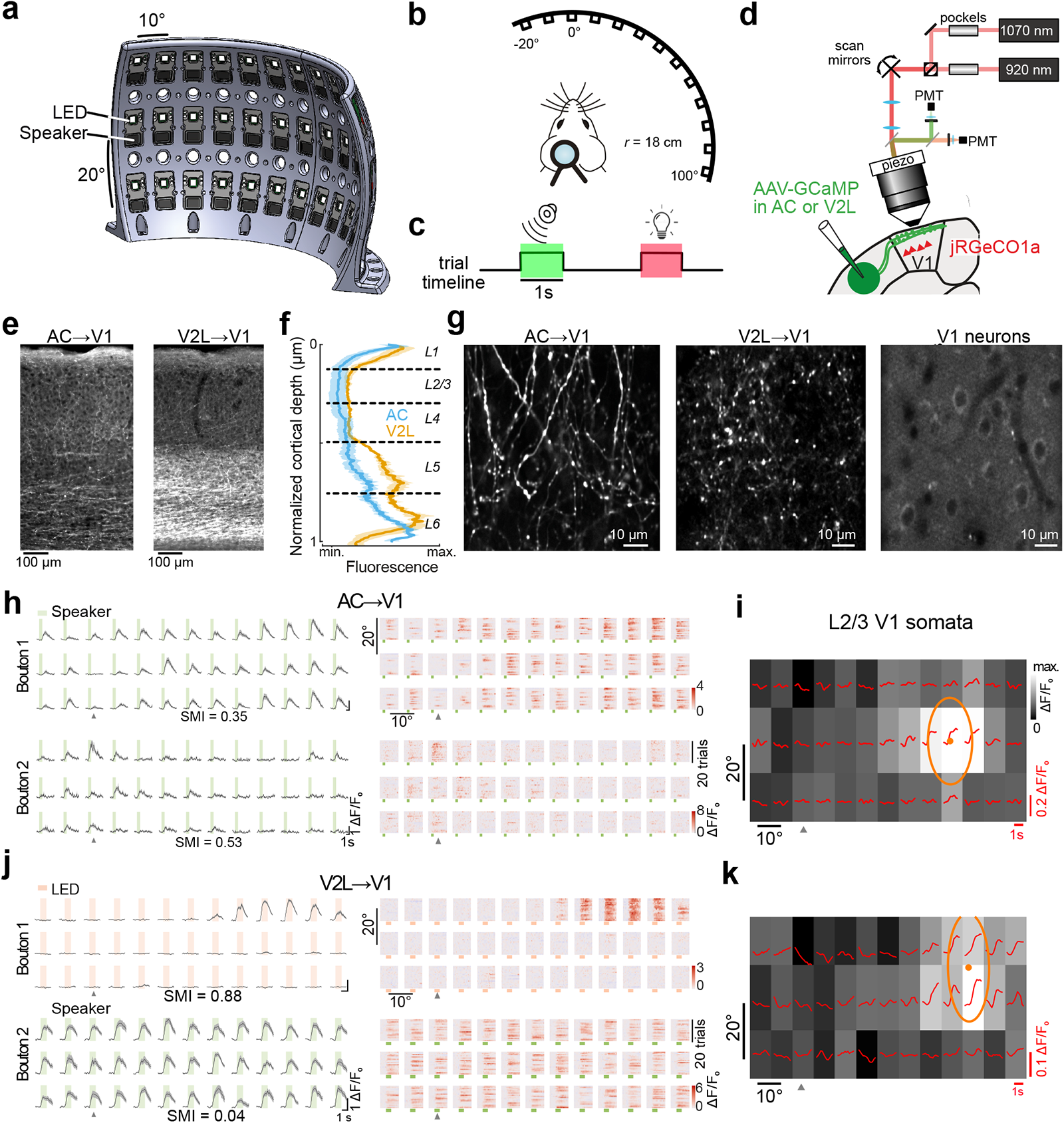
Measuring auditory and visual receptive fields in AC and V2L inputs and how they compare with those of their target V1 neurons. **a** 3D rendering of the LED and loudspeaker array. **b** Spatial relation between the head-fixed mice and the speaker array. **c** Trial structure. **d** Schematic of the dual-color, volumetric two-photon calcium imaging configuration. **e** Coronal histological section of AC (left) and V2L axons (right) in V1. **f** Normalized fluorescence intensity from AC (blue) and V2L axons (yellow) at different cortical depths. Mean (solid line) ± s.e.m. across mice (shaded area); n = 7 AC-injected mice and n = 3 V2L-injected mice. **g** Two-photon field of view in L1 of example GCaMP6s-expressing AC or V2L axons and jRGECO1a-expressing somatas recorded in L2/3 beneath the AC/V2L axons in V1. **h** Left, Responses in example AC boutons from the same field of view in L1 of V1 to sounds from the different speakers in the array. Green shaded area displays sound presentation window. Mean (solid line) ± s.e.m (shaded area). Right, Single trial responses of the same boutons. Sound presentation, green rectangle (1 s). Arrowhead shows midline position. SMI, spatial modulation index. **i** Population visual responses in the jRGECO1a-expressing neurons in L2/3 underneath the boutons in **h**. Average fluorescence signals (red traces) and response over the stimulus presentation window (grayscale color map) from stimuli at the corresponding location in the speaker and LED array. Ellipse, fitted RF; dot, RF center. **j** Visual (top) and auditory (bottom) responses in example V2L boutons in the same field of view in L1 of V1. Salmon shaded area denotes visual stimulus presentation. **k** Population visual responses of L2/3 somata underneath the V2L boutons in **j**.

We measured the spatial specificity of sound evoked activity in AC➔V1 projections using two-photon recordings of axons (Figure 1d). We injected an adeno-associated virus (AAV) encoding the genetically encoded calcium indicator GCaMP6s into two sites along the rostro-caudal extent of the left AC (AAV1-hSyn-GCaMP6s; Supplementary Figure 1a). By using transgenic mice (Thy1-jRGECO1a) ^42^, we simultaneously expressed the red calcium indicator jRGECO1a in V1 neurons and measured their visual RFs using the LED array (Figure 1g,i,k). As neighboring V1 neurons in the small patch of cortex contained within a field of view have similar visual RFs ^43^, this allowed comparing the auditory spatial information in AC➔V1 axons to the visual RF of their postsynaptic neurons, despite being unable to identify them. To determine to what extent auditory RFs were specific to AC➔V1 axons, we compared them against V2L axons ^41^ by injecting GCaMP6s in V2L in an another cohort of mice (Supplementary Figure 1b).

We ensured the accuracy of the injections post hoc by registering coronal histological sections to the brain atlas. We verified that in all the analyzed animals the injection sites were confined to AC or V2L and that the thalamic projections from GCaMP6s expressing neurons mainly targeted the medial geniculate nucleus or the lateroposterior nucleus in AC and V2L-injected mice, respectively (Supplementary Figure 1a-c). AC and V2L projections in V1 mainly innervated layer (L)1 but also sent sparser projections throughout the cortical depth as previously described (Figure 1e,f) ^21, 44, 45^. When imaged *in vivo*, both AC and V2L axons formed a dense mesh in L1 with many *en passant* boutons (Figure 1g).

We extracted fluorescence traces from individual AC or V2L boutons and measured their responses to sounds and flashes of light at different locations. While a significant fraction of AC boutons were responsive to auditory stimulus (fraction responsive, observed vs. time-shuffled data: p = 0.002, two-sided paired t-test, n = 8 mice), the fraction showing light-evoked responses were not significantly larger than the shuffled data (p = 0.08) (Figure 2a,b). In a subset of experiments, we confirmed that AC boutons were frequency-tuned, as expected from auditory responses in AC neurons (Supplementary Figure 2a). In V2L boutons, a large fraction showed visually-evoked responses (p = 0.02, n = 3 mice) that were spatially clustered, reflecting their spatial RF (Figure 1j and Figure 2a,b). In addition, a subpopulation of non-visually responsive V2L boutons responded to auditory stimulation (fraction responsive, observed vs. time-shuffled data: two-sided paired t-test, p = 0.05; Figure 1j and Figure 2a,b). Subsets of AC boutons showed reliable responses to sounds in specific locations.

**Figure 2.**
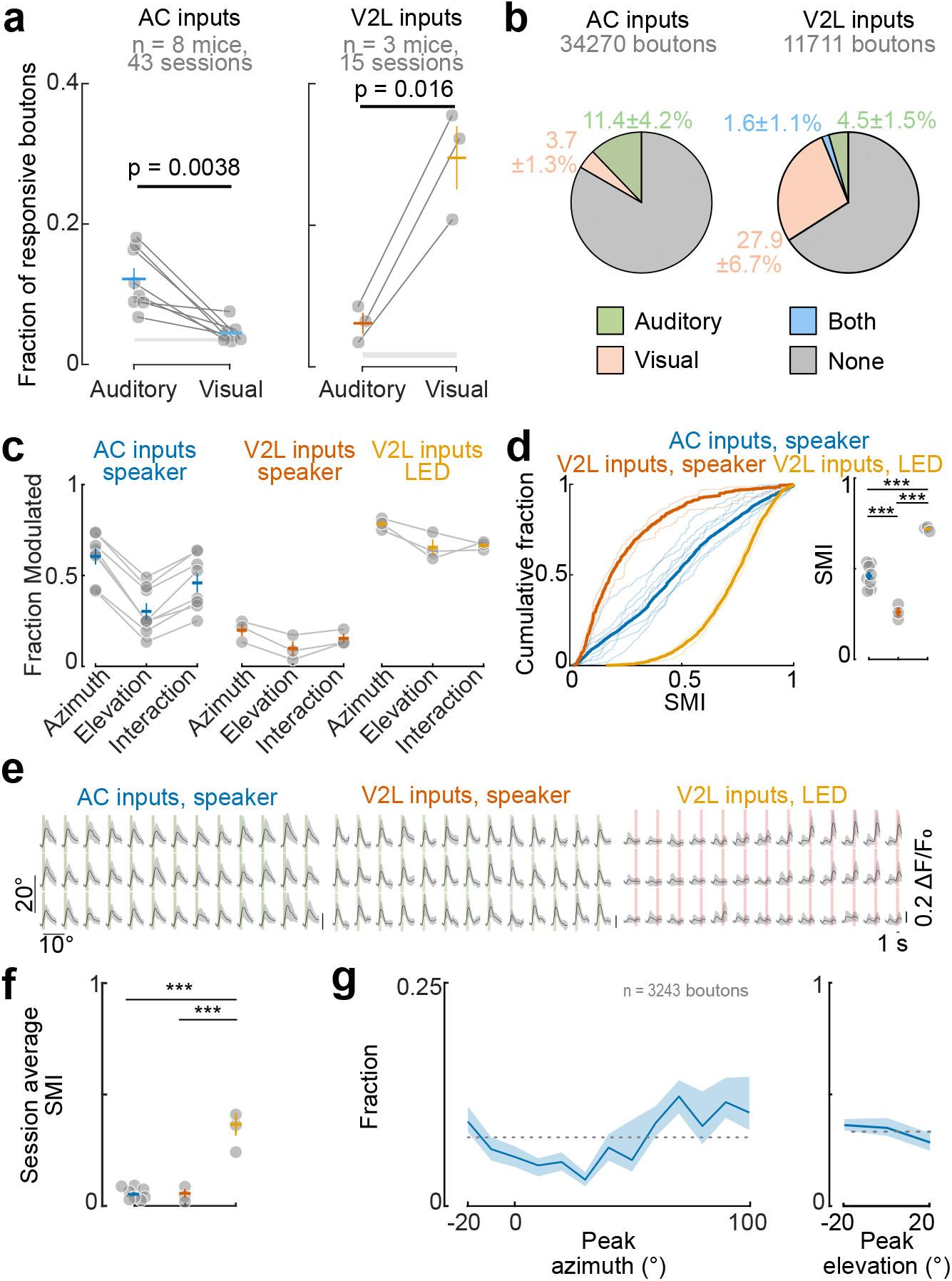
Modality and spatial sensitivity in AC and V2L inputs to V1. **a** Fraction of responsive boutons for sound and visual stimulus in AC and V2L boutons. Gray circles, mean across individual mice; colored crosses, mean ± s.e.m. across mice; gray shaded area, mean fraction of responsive boutons from time-shuffled data ± s.e.m. across mice. Two-sided paired t-test. **b** Fraction of boutons responsive to auditory, visual or both stimuli in AC and V2L axons. Percentages are mean ± s.d. across mice. Fraction of AC boutons responsive to both stimuli was < 1%. **c** Fraction of boutons modulated by sound location in azimuth, elevation or the interaction of the two. Dots, individual mice; crosses, mean ± s.e.m; n = 8 AC-injected and n= 3 V2L-injected mice. **d** Left, Distribution of the spatial modulation index (SMI). 3-sample Anderson-Darling test, p = 0. Thin lines, individual mice; thick lines, average per group. Right, Mean SMI. Dots, mice; cross, mean ± s.e.m. One-way ANOVA: F(2,11)=56.1, p=1.7x10^-6^. Tukey’s post-hoc test: AC, speaker vs. V2L, speaker, p=10^-4^; AC, speaker vs. V2L, LED, p=10^-5^; AC, speaker vs. V2L, LED, p=10^-6^. **e** Example session averages across all responsive boutons to sound and LED flashes at different positions (same sessions as in Figure 1 **h-k**). Green shaded area: sound presentation; Salmon shaded area: visual stimulus presentation. **f** SMI of the average across boutons. Each dot is the average per mouse. Crosses, mean ± s.e.m. One-way ANOVA: F(2,11)=46.5, p=4.3x10^-6^. Tukey’s post-hoc test: AC, speaker vs. V2L, speaker, p=0.97; AC, speaker vs. V2L, LED, p=10^-6^; AC, speaker vs. V2L, LED, p=10^-5^. **g** Distribution of the peak azimuth (left) and elevation (right) across AC axon boutons. Line, mean; shading, 95% confidence interval. Dotted line denotes the uniform distribution.

While auditory responses in AC boutons were often modulated by sound location, those of V2L boutons tended to spatially homogenous (Figure 1h; Figure 2c). Visual responses in V2L boutons were also modulated by LED location, as expected from their known visual RFs ^41^ (Figure 1j; Figure 2c). A larger fraction of the auditory responses in AC boutons was modulated by the azimuth compared to elevation of the sound source (Figure 2c). This difference was not due to the larger angular span in azimuth of the speaker array as it was preserved when restricting the analyses to stimuli from 40° x 40° isotropic arrays (20° step) of 9 speakers (p = 0.027, n = 8 AC mice, two-sided paired t-test; Methods).

In each recording session, we observed a diversity of spatial response profiles in auditory responses in AC boutons, varying from entire or hemifield responses to narrowly tuned responses for different locations (Figure 1h). In many cases, neighboring boutons recorded in the same imaging session showed diverse spatial profiles in their sound-evoked activity (Figure 1h). On the contrary, responses in V2L boutons were less varied. Most auditory-responsive boutons similarly responded to auditory stimulus in all locations, and visually-responsive bouton responded to restricted spatial locations (Figure 1j). To quantify the extent to which bouton responses were modulated by space, we calculated the spatial modulation index (SMI) of each bouton (Methods). The SMI has a value of 1 when responses are confined to a single location and 0 when their amplitude is equal across all locations (see SMI for example boutons in Figure 1h,j). The SMI was largest for visual responses in V2L boutons, as expected from the known spatial selectivity of visual cortex neurons (Figure 2d). The auditory responses of AC➔V1 boutons were, on average, more spatially selective than the auditory responses of V2L boutons but less selective than their visual responses (Figure 2d).

We measured the extent of the locations from which responses could be elicited in AC and V2L inputs to V1 at each recording site by averaging all the responsive boutons and calculating the SMI (Figure 2e,f). As expected, the averaged visual response across V2L boutons was restricted to specific LEDs, as expected from the retinotopic specificity of this projection ^41^. On the contrary, average auditory responses in both AC and V2L inputs were widely spread across all the speaker locations. Accordingly, the SMI of the average response per imaged location was lower for auditory responses in both projection types when compared with the visual responses of V2L boutons (Figure 2f).

The position eliciting the largest response varied across AC axons, with a slight enhancement in boutons tuned to high-azimuth positions (Fig 2g). As for visual responses in V2L boutons, sound-evoked responses in AC boutons decreased monotonically with increasing distance from the boutons’ preferred location (Supplementary Figure 2b).

The previous observations showed that AC➔V1 inputs have spatially confined auditory RFs that are on average broader than the visual RFs of V2L➔V1 inputs. We assessed whether the spatial information conveyed by AC➔V1 inputs is sufficient to decode the position of the stimulus on a single trial basis using a naïve Bayesian decoder (Methods). We grouped highly correlated boutons into one functional unit as they are likely to belong to the same axon ^46, 47^ and measured the ability of the decoder to accurately locate sounds on a per trial basis using all the responsive axons from one imaging session. The distribution of the likelihoods across stimulus locations obtained by the decoder displayed a spatial component, with decaying likelihood away from the actual stimulus position (Figure 3a). Consistently, the distribution of the azimuthal distance between predicted and actual stimulus location was skewed towards small distances while larger distances were less often predicted than chance (Supplementary Figure 3c). We scored the accuracy of the decoder by measuring the fraction of times that the decoded position with the largest likelihood was within 10° of the actual one. Using AC auditory responses, speaker position could be decoded better than that of the axon identity-shuffled data in both elevation and azimuthal space (Figure 3b). Decoding performance did not depend on the position of the speaker (Supplementary Figure 3a), nor on the retinotopic position in V1 where the axons were recorded from (Supplementary Figure 3b). In contrast, decoding accuracy from auditory responses in V2L axons was not significantly greater than the shuffle control (Figure 3b; Supplementary Figure 3a,c). The location of visual stimuli, however, could be accurately estimated from V2L axons, as expected from their spatially confined visual RF (Figure 3b; Supplementary Figure 3a,c). We compared the distance between the actual and predicted stimulus location as a function of the number of axons sampled across projection types. Spatial accuracy using auditory responses in AC axons was greater compared to V2L axons but lower than when using visual responses in V2L axons (Figure 3c).

**Figure 3.**
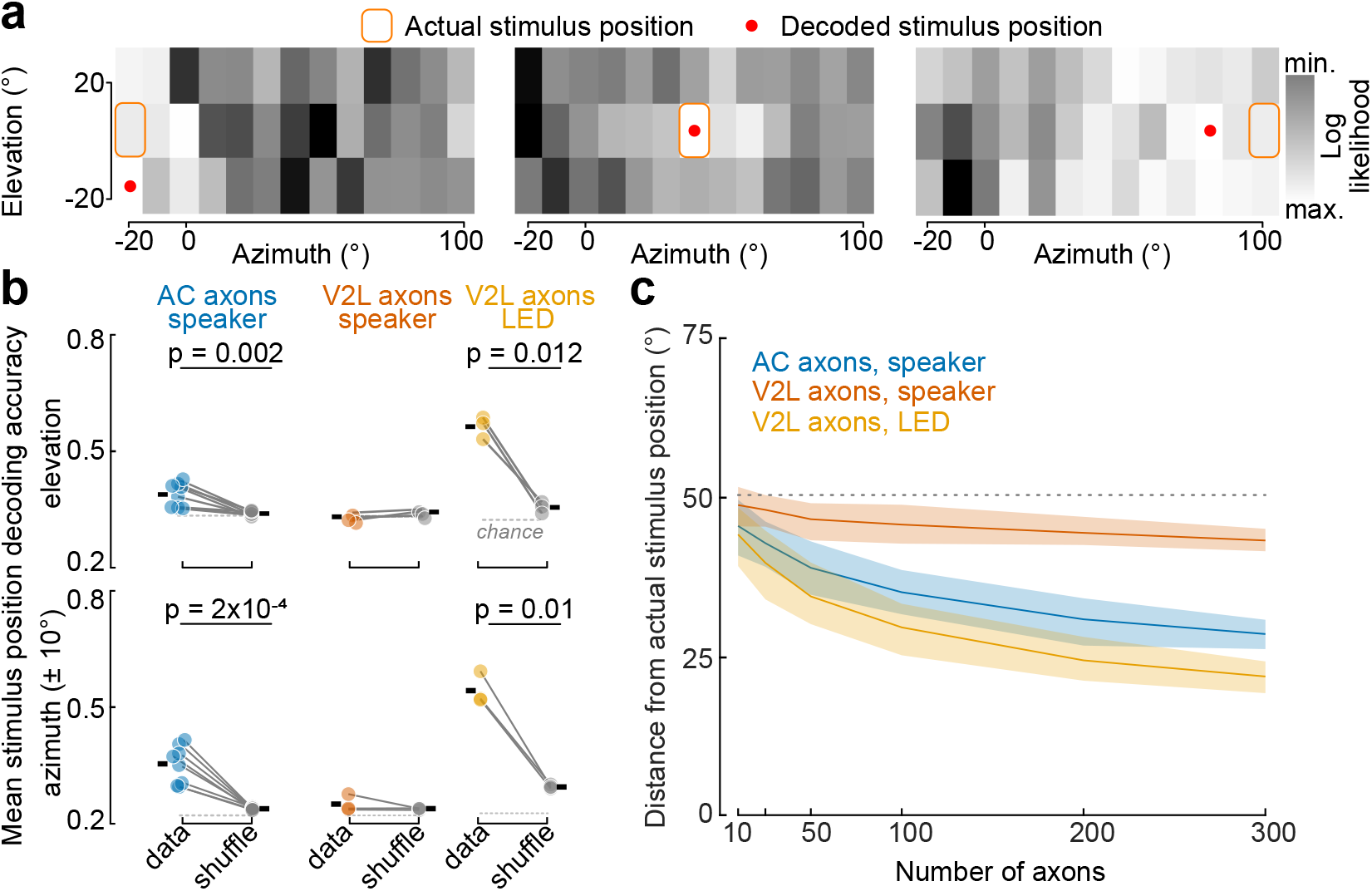
Sound location can be decoded from AC inputs to V1. **a** Log likelihood distribution across speaker positions for three example trials. Circle, maximum of the likelihood; square, actual speaker used in that trial. **b** Elevation and azimuth stimulus position decoding accuracy. Colored dots, individual mice (average across imaging sessions); grey dots, shuffled data; ticks, mean across mice. Two-sided paired t-test: n = 8 AC-injected mice, 42 sessions; n=3 V2L-injected mice, 15 sessions. **c** Distance between decoded and actual stimulus location as a function of number of axons. Solid line, mean; shading, 95% confidence interval; dotted line, chance level.

We conclude that AC axons, but not those from V2L, relay signals about the location of sounds to V1. The combined inputs from many AC boutons in L1 are sufficient for inferring the location of sounds but with lower accuracy than when locating visual stimuli using V2L boutons.

### Spatial auditory responses in AC axons are independent of behavioral responses

Auditory modulations of neural activity in V1 neurons is thought to be driven, at least in part, by sound-evoked changes in internal state and uninstructed body movements ^14, 29, 30^. However, the location-specific sound-evoked activity we observed in AC axons is not consistent with the known features of behaviorally driven cortical modulations. First, these modulations are low dimensional ^30, 48^, i.e. they affect all neurons similarly, while the spatial profile of the sound evoked activity in AC➔V1 inputs was diverse across boutons, even within the same session (Figure 2a). Second, movement-induced modulations of neuronal activity are broadly distributed across brain structures ^48–51^, which contrasts with the fact that sound-evoked spatially restricted responses were present in AC, but largely absent in V2L inputs to V1 (Figure 2; Figure 3). To further confirm that spatial RF in AC boutons were independent of uninstructed movements, we analyzed the behavioral responses to sound from the different locations by measuring pupil dilation (5 AC mice and the 3 V2L mice) in a subset of the experiments. Sound did indeed induce pupil dilation (Supplementary Figure 5a), but it was indifferent, on average, to speaker locations (Supplementary Figure 5b), Furthermore, speaker location could not be decoded from the pupil data (Supplementary Figure 5c). We also reached the same conclusion when analyzing face movements, as these had been shown to better predict auditory-driven neuronal activity ^30^(Supplementary Figure 5e). Finally, sound-evoked responses in AC➔V1 boutons were poorly correlated with pupil size. In contrast, pupil size was more correlated with auditory responses in V2L than in AC boutons (Supplementary Figure 5d). Thus, while the sound evoked responses in V2L boutons are consistent with behavioral modulations, their spatially pattern in AC➔V1 boutons is independent of them.

### Auditory spatial information in AC axons to V1 is not topographic

We then investigated whether the auditory spatial RFs of AC➔V1 inputs were topographically organized with respect to the V1 retinotopic map. We measured the azimuthal position of the speaker that reliably elicited the strongest response for each AC bouton (best azimuth) and calculated the mean across boutons per session. We then compared the mean azimuth peak of the AC boutons with the azimuth center of the visual RF of the population of jRGeCO1a-expressing L2/3 V1 somatas below them (Figure 1i,k).

The auditory RFs of AC boutons were not topographically organized in V1 as the average best azimuth bore no relation with the V1 neurons below them (Figure 4a). In contrast, the visual RFs of the V2L boutons were topographically organized and matched with those of the V1 neurons they target, as previously described (Figure 4b) ^41^.

**Figure 4.**
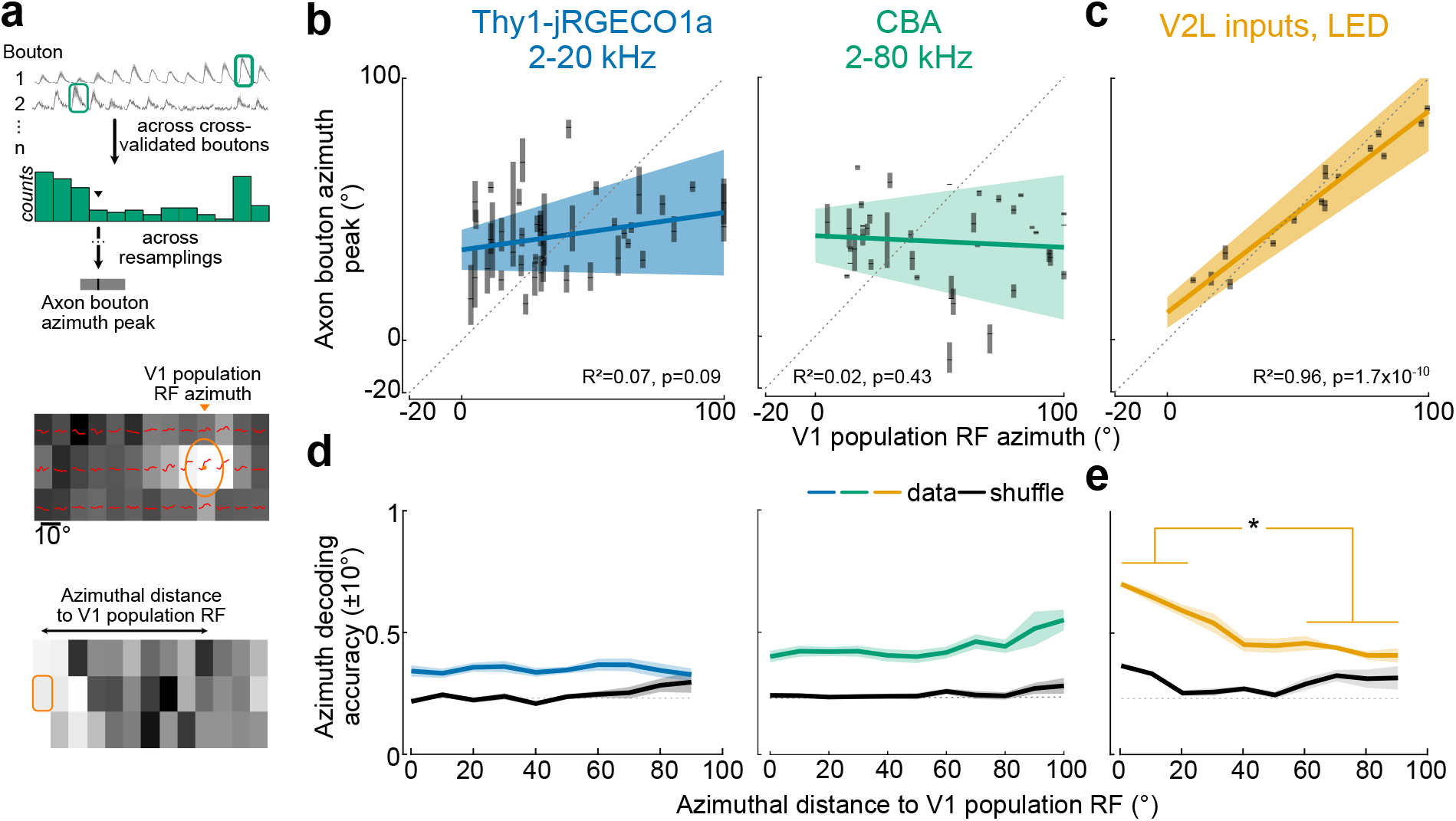
AC inputs in V1 are not retinotopically matched. **a** Top, Example bouton responses, averaged across elevation. Same boutons as in Figure 1h. Green boxes denote azimuth peak. Median of the distribution (arrowhead) is used to estimate the population azimuth peak. This is repeated over 100 iterations of resampling to estimate the mean and 95% confidence interval (black bar and gray shaded area) of the axon bouton population peak response in azimuth per imaging session. Middle, V1 somata population RF azimuth (orange arrowhead) (heatmap as in Figure 1i). Bottom: Decoding of an example trial at -20° in azimuth (orange box, as in Figure 3a). The distance between the stimulus position and V1 population RF azimuth is represented on the abscissa in **d,e**. **b** Mean peak azimuth of the sound evoked response across AC boutons as a function of the population RF center of V1 neurons for each imaging session. Left, Thy1-jRGECO1a mice, 2-20 kHz auditory white noise; n = 39 imaging sessions, 8 mice. Right, CBA mice, 2-80 kHz auditory white noise; n = 47 imaging sessions, 6 mice. Ticks, median; grey shading, 95% confidence interval. Colored lines and values correspond to the linear regression of the mean values; colored shading, 95% confidence interval. Dashed lines, identity lines. **c** Peak azimuth of visual response in V2L boutons; n = 15 imaging sessions, 3 mice. **d** Decoding accuracy of sound azimuth using AC axons as a function of the distance between speaker location and V1 population RF center. Left, AC axons of Thy1-jRGECO1a mice, 2-20 kHz auditory white noise. Right, AC axons of CBA mice, 2-80 kHz auditory white noise. Data is mean ± s.e.m. across mice. Dotted line, chance. One-way ANOVA, Thy1-jRGECO1a: F(9,69) = 0.25, p = 0.98; n = 8 mice, 38 imaging sessions; CBA: F(10,40) = 1.92, p = 0.07, n = 5 mice, 24 imaging sessions. **e** Decoding accuracy of light stimulus azimuth using V2L axons as a function of the distance between the LED location and V1 population RF center. One-way ANOVA: F(10,22) = 15.2, p = 1.1x10^-7^; N = 3, n=12. *: p < 0.05 for all paired comparison with Tukey post-hoc test.

We checked if the location of sounds could be decoded differently from AC➔V1 inputs when they correspond to the retinotopically-matched location in V1, despite the lack of retinotopic organization in the projection. We found that the decoder’s performance was similar regardless of the speaker’s location relative to the V1 azimuthal position where AC inputs were recorded (Figure 4b), indicating no functional selective mapping of auditory space to V1. In contrast, decoding performance increased with increased proximity to V1 RF center in visual responses in LM➔V1 axons, consistent with the retinotopic alignment of these inputs (Figure 4d).

### The absence of topographic organization in the auditory spatial responses of AC**➔**V1 axons is not due to a lack of high-frequency cues

Recent work has shown that high-frequency cues (>20 kHz) are necessary for the topographic alignment of the auditory and visual RF in azimuths in the SC ^10^. As a consequence, C57BL/6 mice do not show a topographic auditory and visual alignment in the SC due to a selective early onset high-frequency hearing deficit ^52^. Because Thy1-jRGECO1a mice were on a C57BL/6 background ^42^, they might be susceptible to high-frequency hearing deficit as well. Therefore, we repeated our experiments using broadband white noise that includes high frequencies (2-80 kHz) in CBA mice, as this strain does not suffer age-related high-frequency hearing loss ^53^, and in another group of Thy1-jRGECO1a mice.

We used AAV injections to transfect AC➔V1 inputs with GCaMP8m ^54^ and V1 with jRGECO1a (in CBA mice). As before, AC➔V1 boutons in both groups showed a rich variety of location specific sound evoked responses. Consistent with their role in sound localization, in both CBA and Thy1-jRGECO1a mice, the decoding accuracy of speaker azimuthal location was higher than that of the shuffled data (Supplementary Figure 5a) and more precise than when using band-limited sounds in Thy1-jRGECO1a mice (Supplementary Figure 5b). As before, however, we did not observe a correlation between the auditory RF azimuthal centers and the visual population RF azimuthal center from V1 L2/3 somatas (Figure 4a, Supplementary Figure 5d). Similarly, decoding accuracy did not depend on the retinotopic position in V1 of the AC➔V1 inputs (Figure 4c, Supplementary Figure 5c). Thus, while the presence of high-frequency cues enhances the location-specific information available in AC➔V1 inputs, the lack of a topographic organization we observed is not due to a deficiency of these cues during development or during our measurements.

### AC**➔**V1 axons represent the location of sounds in both the ipsi- and contra-lateral hemifields

Many AC boutons in V1 exhibited peak responses when sound was presented in one of the two ipsilateral locations in our speaker array. (Figure 2f). To measure to what extent AC inputs represent spatial locations outside the contralateral visual field of V1, we mounted a single loudspeaker on a rotating arm (Figure 5a) and presented sounds over 180° around the head at a single elevation, spanning 90° ipsi- and contra-laterally. Consistent with the measurements using a speaker array, many AC➔V1 axons were spatially tuned. We confirmed that the spatially specific sound responses obtained with the speaker array were similar to those measured using the rotating speaker (Supplementary Figure 6).

**Figure 5.**
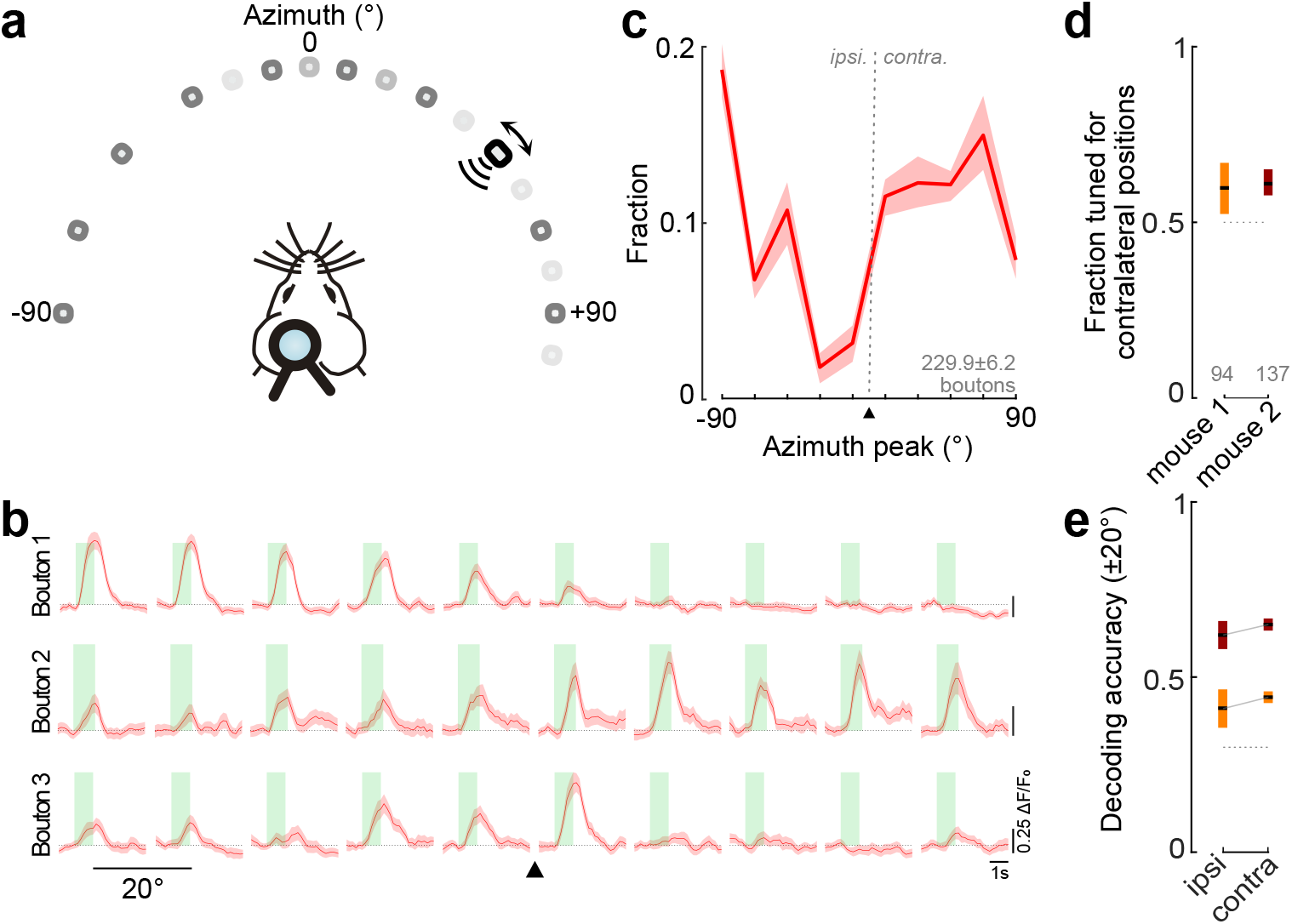
AC axons to V1 L1 are tuned to sounds located in both ipsi- and contra-lateral hemifields. **a** A single loudspeaker (black rectangle, 2-80 kHz white noise) was moved between -90° to +90° in azimuth, 20° spacing (dark gray positions). Light gray positions represent positions sampled during the same experiment, analyzed separately (see **Supplementary** Figure 6). **b** Three example AC bouton responses to sounds played at different ipsi- and contra-lateral azimuthal positions. Solid lines, mean response; shaded area, s.e.m. Arrowhead, midline. **c** Distribution of azimuth peak across boutons. Solid line, mean; shaded area, 95% confidence interval; number of cross-validated boutons across resampling iterations: n = 229 ± 6.2 (mean ± s.d.) from 2 mice. Arrowhead, midline (azimuth = 0°). **d** Fraction of contralateraly tuned boutons for 2 mice. Ticks, mean across resampling iterations; shaded areas, 95% confidence interval; Gray: mean number of cross-validated boutons. **e** Decoding performance averaged across sound located at ipsi- and contra-lateral positions (mean ± s.e.m across sessions per mice, same colors as in **d**; dotted line, mean using shuffled data.

Neighboring boutons showed different spatial RFs that, in many cases, peaked in different hemifields (Figure 5b). Boutons tuned to contralateral and ipsilateral locations were both abundant, with a only slight, but significant, overrepresentation of contralaterally-tuned ones (Figure 5c,d). Consequently, the locations of sounds in both hemifields could be similarly decoded (Figure 5e). Thus, many AC➔V1 inputs relay localization signals about sounds originating in regions outside the visual hemifield represented in V1.

### Audio-visual interactions in V1 neurons do not depend on the spatial coherence of the stimuli

We checked if V1 responses are influenced by the spatial congruence of auditory and visual stimuli, even though the inputs to V1 from the AC were not retinotopically matched. We recorded audio-visual responses from V1 L2/3 neurons using two-photon calcium imaging in mice constitutively expressing GCaMP6f in pyramidal neurons (Figure 6a).

**Figure 6.**
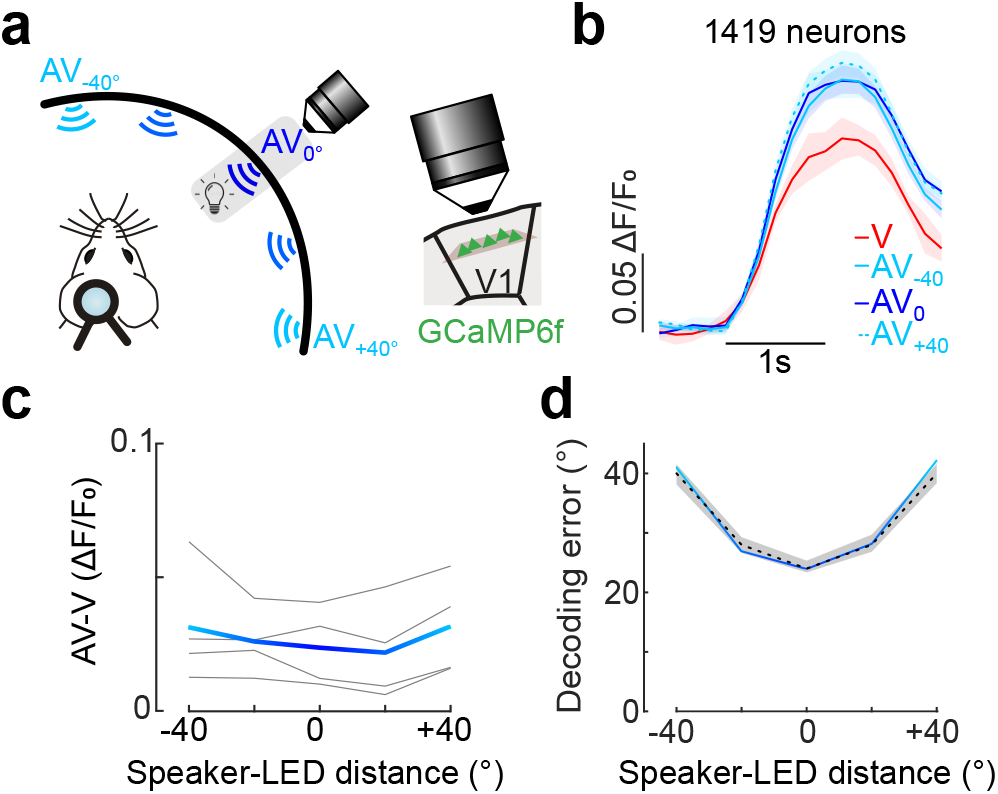
Audiovisual interactions in V1 do not depend on the spatial coherence between the two stimuli. **a** Imaging of GCaMP6f-expressing L2/3 neurons in V1 during audiovisual experiments. An LED located within the RFs of the recorded neurons (gray shaded area; azimuth RF center range: 40-60°) was flashed while an auditory stimulus was presented at one of the azimuthal locations (-40° to 40°, 20° steps). 0° denotes matched speaker and LED position, negative and positive values correspond to speakers in lower and higher azimuthal positions, respectively. Audiovisual interactions were tested across different loudness and brightness levels. **b** Average visual (V) and audiovisual (AV) responses for 3 example speaker positions for all the recorded neurons of an example mouse, averaged across brightness and loudness levels. The relative position of the LED and speakers of each stimuli is illustrated in panel **a**. Data are mean across neurons ± s.e.m. Visual responses in V1 neurons were enhanced by simultaneous auditory stimulation (two-sided paired t-test, p = 6 x 10^-6^). Black bar represents stimulation window (1 s). **c** Distance between LED and speaker did not influence the strength of the AV modulation (averaged across loudnesses and brightnesses for visualization purposes). Three-way repeated measure ANOVA, effect of position: F(6,18) = 2.14, p = 0.098; n = 4 mice. Gray lines, average across cells per mice; colored line, average across mice, with distance between speaker and LED color-coded. **d** Decoding error (distance between actual and decoded position) was similar to that of the shuffle data for all speaker-LED distances. Colored line, average across mice, with distance between speaker and LED color-coded; gray shaded area, 95% confidence interval from shuffle data; dotted line, chance level.

We presented stimuli at different positions using the LED and speaker array. We presented light flashes within the RF of the recorded neurons either alone (V) or concurrently with white noise sound bursts (AV) from a speaker at the same spatial location (0°) or ± 20° and ± 40° away in azimuth. Auditory responses were not significantly different across positions (One-way ANOVA: F(6)=0.61, p = 0.72, n = 4 mice). We tested different combinations of LED brightness and speaker loudness. Visual responses in L2/3 V1 neurons were on average larger in the presence of sound (Figure 6b; two-sided paired t-test, p = 10^-7^) ^14, 17, 29^. However, the magnitude of the AV enhancement did not depend on the distance of the auditory and visual stimuli (Figure 6c), or their brightness or loudness (Three-way repeated measure ANOVA: effect of brightness, F(2,6) = 0.76, p = 0.5; effect of loudness, F(2,6) = 3.01, p = 0.12; n = 4 mice). Consistent with this, the speaker location could not be decoded from the AV responses of V1 neurons (Figure 6d).

In a subset of experiments, we checked if spatially matched and non-matched AV responses in V1 would be different in cases when the two stimuli were further separated. As before, AV responses were indistinguishable when the speaker was separated from the LED by 80° in azimuth and they were spatially matched (two-sided paired t-test: p = 0.44, n = 5 mice).

We conclude that in passively stimulated mice sound modulations of visual responses in V1 do not depend on the spatial register of the stimuli from the two modalities.

## DISCUSSION

We measured the relay of auditory and visual information from the AC to V1. Our findings show that while many AC inputs had broad spatial sound-evoked responses, others responded to sounds at specific ipsi- or contralateral locations. Through decoding analysis, we demonstrated that by sampling enough AC inputs in L1, V1 neurons could accurately estimate the location of sounds. Unlike visual RFs from other visual areas, the auditory RFs of AC inputs to V1 were non-topographic and did not bear any relation to the retinotopic map in V1. Consistent with the lack of topographic projection, we demonstrated that modulations of visually evoked responses in V1 did not depend on the spatial relationship of auditory and visual stimuli.

### AC relays information about the location of sounds to V1

We found auditory responses were much more frequent than visual ones in the projection. Thus, in passively stimulated animals, unlike in mice that learned an audio-visual association^44^, AC inputs to V1 are dominated by auditory responses over visual ones. These responses were independent of any uninstructed body movements and were consistent with somatic recording from V1-projecting AC neurons ^17^, confirming that auditory responses are present in AC inputs to L1 of V1.

As in the AC as a whole ^32, 34, 35, 40, 55–57^, AC afferents were spatially non-selective, or selective to varying degrees to sounds in specific psi- or contralateral locations. Given their broader spatial RFs, single AC boutons have limited information about the location of sounds. As a population, however, they have the capability to provide an accurate estimate of the location of sounds to postsynaptic V1 neurons. Thus, AC inputs in V1 provide a rich neural substrate from binding spatially and temporarily congruent AV stimuli.

### Spatial information in AC inputs to V1 is not topographic

While many AC-+V1 inputs had spatially confined auditory RFs, these differed from the visual RF of the V1 neurons they innervated in several aspects:

1) AC inputs to V1 were less spatially specific than the visual RFs of V1 neurons (Figure 3 and Figure 4), consistent with somatic recordings from AC neurons ^32, 58^,. 2) In addition, the total span of the auditory RFs in AC inputs to V1 was larger than the visual field coverage of V1. That is, while within each hemisphere, mouse V1 mainly represents the contralateral visual space and only a small portion of ipsilateral frontally facing space, a large fraction of AC inputs was selective for ipsilaterally sounds originating outside this range (Figure 5). Thus, AC afferents in V1 relay information about sounds located outside the field of view of the targeted area. 3) We did not find any evidence of a topographic organization in the auditory spatial information relayed by AC to V1. This was true even in experiments with sound stimulus covering the full auditory spectrum (2-80 kHz white noise and in mice that do not suffer from age-related high-frequency hearing loss ^53, 59^. While AC is tonotopically organized, it harbors no map of auditory space across species ^40^, including mice ^32^. Our observations show that even when innervating an area with an accurate spatial map as V1, AC inputs remain non-topographically organized. Consistent with this absence of retinotopic organization, sound-induced modulations of visual responses did not depend on the spatial distance between the two stimuli (Figure 6). This contrasts with superior and inferior colliculi, were auditory and visual maps are aligned and responses are specifically enhanced by spatially congruent bimodal stimulus ^10–12, 60, 61^. Thus, even though neurons in the two cortical areas share a common representational axis, i.e., they both represent the location of the stimuli, the connections between them do not follow a like-to-like connectivity. This contrasts with projections from higher-order visual and frontal areas to V1 that are topographically organized to match the retinotopy of V1, e.g. the visual RF of the afferent inputs are on average matched with those of their targeted V1 neurons (Figure 5b) ^25, 41, 62, 63^. This difference in functional connectivity might be due to several features. On one hand, unlike AC inputs, higher-order visual and frontal areas convey spatial information from the same modality to V1. Sound location is encoded in head-centered coordinates, while light source location is encoded in eye-centered coordinates. Our experiments were performed in head-fixed mice, yet pinna position affects the relation of spectral cues to sound location. Thus, variations in both ear and eye positions might preclude the precise alignment of visual and auditory spatial maps. Yet, in the SC of mice, auditory and visual maps are aligned ^10^, showing that audiovisual spatial correspondence is possible despite the different coordinate frames.

On the other hand, AC inputs to V1 are not reciprocated, unlike those from higher-order visual and frontal areas, AC being devoid of V1 inputs in mice and other species ^28, 64^. Thus, retinotopically matched interactions across cortical projections might require recurrent interactions that are present in many cortical feedback inputs to V1 ^45, 65, 66^ but are absent in the AC➔V1 projection.

### Implications for the functional role of AC afferents in V1

What could the role of the non-topographic signals encoding sound location that are relayed from AC to V1 be? The organization of direct AC➔V1 cortico-cortical inputs and the independence of AV modulations on the spatial coherence of the two stimuli differs with the spatial register found the SC and the inferior colliculus^1, 10, 60^. These differences suggest that audiovisual interactions mediated by direct cortico-cortical connections might play a fundamentally different role than in midbrain structures.

Direct, unidirectional projections from AC to V1 are present in many species, suggesting a conserved function. Bimodal stimuli that are congruent in space and time result in perceptual and reaction time improvements in various tasks ^1^. However the role played by projections directly linking sensory areas of the neocortex remain unclear ^13^. Given that auditory cortical responses are faster than visual ones, one possibility is that direct AC inputs sensitize V1 neurons with RFs overlapping with salient audiovisual objects, drawing attention to them and facilitating their discrimination ^67, 68^. Consistent with this, visual stimuli are perceptually more salient when they are in spatial register with the auditory ones ^69–71^. However, such a mechanism would require aligned retinotopically matched AC afferents in V1. In addition, this hypothesis leaves inputs from AC neurons with RFs located outside the span of the innervated V1, such as from laterally located ipsilateral positions, without a function. One possibility is that V1 neurons integrate sound localization signals from AC inputs in L1 with upcoming motor commands for oriented body and head movements relayed from other sources. By integrating motor commands, V1 neurons could re-reference auditory localization signals in AC axons to retinotopic coordinates and become sensitized to the visual stimuli the AC inputs anticipate ^67^. Such a mechanism would require spatially specific sound signals in AC inputs, but these do not have to be necessarily retinotopically matched, as observed here.

Another alternative would be that, while integrating sound information with visual inputs, V1 is not involved in detecting of low-level features, such as the location of sounds, but uses the diverse and distributed spatial information available in AC➔V1 inputs for high-order analyses of auditory space, such as the relative location of different sound sources for integration with visual information. Future studies addressing how sound location-dependent visual modulations depend on experience will provide insights on how V1 neurons integrate the rich auditory spatial signals available in L1.

## Acknowledgements

We thank Matthijs Oude Lohuis, Alfonso Renart and all members of the Petreanu Laboratory for comments on the work, the Champalimaud Research Hardware Platform for technical support, and Adrien Jouary and Davide Reato for help with experimental procedures and data analysis. This work was supported by a Marie Skłodowska-Curie individual fellowships (MSCA 798941; C.M.) and Human Frontiers Scientific Program fellowship (LT000064/2018-L; C.M.). L.P was supported by grants from Fundação para a Ciência e a Tecnologia (PTDC/MED449 NEU/30328/2017 and PTDC/MED-NEU/6645/2020), La “Caixa” Foundation (project codes LCF/PR/HR17/52150005 and HR22-00778) and by the Champalimaud Foundation. This work was supported by the research infrastructure CONGENTO, co-financed by Lisboa Regional Operational Programme (Lisboa2020), under the PORTUGAL 2020 Partnership Agreement, through the European Regional Development Fund (ERDF) and Fundação para a Ciência e Tecnologia (Portugal) under the project LISBOA-01-0145-FEDER-022170.

## Author contributions

C.M. and L.P. conceived the study. C.M. performed the experiments and data analysis. C.M. and M.B. performed surgeries and histology. C.M. and L.P. wrote the manuscript.

## Data, software, and hardware availability

Data and custom MATLAB and Bonsai codes will be made available upon publication of the article. The speaker and LED device was built in-house and is open-source and will made available through our institute scientific hardware platform (Champalimaud Hardware Platform; http://www.cf-hw.org).

## METHODS

### Animals

Thy1-jRGECO1a (Tg(Thy1-jRGECO1a)GP8.20Dkim/J, MGI:J:268005, JAX stock #030525), Ai148D (Ai148(TIT2L-GC6f-ICL-tTA2)-D, JAX stock #030328), CBA/CaCrl (Charles River Laboratories, stock #609) and Slc17a7-IRES2-Cre (or Vglut1-IRES2-Cre-D, JAX stock #023527) male and female mice were used in this study. All animal procedures were reviewed by the Champalimaud Centre for the Unknown Ethics Committee guidelines and performed in accordance with the Portuguese Direção Geral de Veterinária.

### Surgeries

Surgeries were performed on young adult mice (8-9 weeks old) under isoflurane anesthesia (1.5%). Bupivacaine (0.05%; injected under the scalp) and buprenorphine (100 μl, 0.1 mg/kg, subcutaneously) provided local and general analgesia, dexamethasone (100 μl, 2 mg/kg, subcutaneously) and sodic cefovecin (100 μl/10 g, 6 mg/kg, subcutaneously) were used to minimize inflammation and infection, respectively. Eyes were protected and kept moist using ophthalmic ointment (Vitaminoftalmina A, Labesfal). Glass injection pipettes (Drummond) were beveled at 45° with a 13-18 µm inner diameter opening and backfilled with mineral oil. A fitted plunger controlled by a hydraulic manipulator (Narashige, MO10) was inserted into the pipette and used to load and inject the viral solution. The skull was thinned above the auditory cortex and the glass injection pipette was lowered through the remaining bone down to the auditory cortex. We injected 50-100 nL of AAV (AAV2/1-Syn-jGCaMP8m-WPRE, Addgene #162375 or AAV2/1-Syn--GCaMP6s-WPRE-SV40, Addgene #100843) either in the AC in two locations (Coordinates, posterior and lateral from Bregma, depth from brain surface: 2.5 mm, 4.3 mm, 0.8 and 1.2 mm depth and 3.0 mm, 4.7 mm and 0.5 mm depth), or in V2L (3.55 mm, 3.65 mm, 0.25 and 0.55 mm depth, 50 nL total). We used either the Thy1-jRGECO1a or CBA mouse strain for AC-injected mice. Two V2L-injected mice were Thy1-RGECO1a strain, and one was C57BL/6 strain. In this mouse and in CBA mice, we additionally injected an AAV encoding for jRGECO1a (AAV2/1-Syn-NES-jRGECO1a-WPRE-SV40, Addgene 100854-AAV1) in 3 locations in V1 (roughly the vertices of a 550 μm-wide equilateral triangle centered 3.5 mm posterior and 2.6 mm lateral to Bregma, 60 nL per location, 0.3 mm depth). AC and V1 coordinates in CBA mice were displaced posteriorly by 150 µm to compensate for the larger size of the brain relative to the Thy1-jRGECO1a mice (estimated from the bregma-to-lambda distance).

To prevent backflow the pipette was kept in the brain for over 5 min after each viral injection. A circular craniotomy was performed over the left visual cortex (diameter: 3 mm; center: 3.5 mm posterior and 2.6 mm lateral to Bregma). The dura was left intact. An imaging window was constructed from three layers of microscope cover glass (1 x 5 mm, 2 x 3 mm diameter, Fisher Scientific, no. 1) joined with a UV-curable optical glue (NOR-61, Norland). The window was placed into the craniotomy and secured in place using black dental cement. A stainless steel headpost, specially designed to allow head fixation to the recording rig without touching the ears, was implanted to the skull with dental acrylic.

### Intrinsic signal imaging

One to two weeks after surgery and before starting two-photon imaging, Intrinsic signal imaging was used to obtain retinotopic maps of the primary visual cortex and surrounding visual areas. Mice were headfixed and lightly anesthetized with isoflurane (1%) and injected intramuscularly with chlorprothixene (1 mg/kg). The eyes were coated with a thin layer of silicone oil (Sigma-Aldrich) to ensure optical clarity during visual stimulation. Optical images of cortical intrinsic signals were recorded using a Retiga QIClick camera (QImaging) controlled using Ephus ^72^ with a high magnification zoom lens (Thorlabs) focused on the brain surface under the glass window at 5 Hz. To measure intrinsic hemodynamic responses, the surface of the cortex was illuminated with a 620-nm red LED while drifting bar stimuli were presented to the right eye in a monitor. An image of the cortical vasculature under the window was obtained using a 535-nm green LED. Azimuth and elevation maps of the visual cortical areas were obtained by calculating the phase for each pixel of the discrete Fourier transform at the visual stimulation frequency ^41, 73^. The hemodynamic delay was canceled by subtracting the phase maps of the experiments from opposing moving stimuli.

### Two-photon imaging

Imaging was performed from 2.5 until 5 weeks post injection. All mice were imaged at < 3.5 months of age, before the development of substantial high-frequency hearing loss ^53, 74^. We used a custom microscope (based on the MIMMS design, Janelia Research Campus, https://www.janelia.org/open-science/mimms) equipped with a resonant scanner, GaAsP photomultiplier tubes (10770PB-40, Hamamatsu) and a 16× (0.8 NA) objective (Nikon), controlled by ScanImage (Vidrio Technologies). O-rings were glued to the headpost, concentric with the center of the cranial window, to form a well for imaging. To prevent stray light from the LEDs or monitor from entering the imaging well, a flexible, conical light shield was attached to the headpost and secured around the imaging objective. GCaMP was excited using a Ti:sapphire laser (Chameleon Ultra II, Coherent) tuned to 920 nm. jRGECO1a was excited using a laser at 1070 nm (Fidelity, Coherent). We performed multi-plane imaging scanning in the axial direction using a piezo actuator (Physik Instrumente) to move the objective. For dual color recordings, we recorded from GCaMP-expressing axons near-simultaneously at two depths in L1, and jRGECO1a at two depths in L2/3 of V1 (field of view [FOV] of 80 × 80 µm, 512 × 512 pixels scanned at ∼6 Hz, ∼20-140 µm deep, 40 µm between planes). Alternatively, 2 planes (∼160-200 µm deep, 40 µm apart) in V1 L2/3 neurons were first collected at ∼10 Hz to map V1 neurons’ RF before performing the GCaMP recording in L1 (4 planes at ∼6 Hz, ∼20-80 µm deep, 20 µm steps). To minimize cross talk between the different calcium indicators when doing quasi simultaneous recordings of GCaMP axons and L2/3 somata ^75^, the 1070 nm beam was turned off (<1 mW of power) when scanning in L1 and the 920 nm one was turned off when scanning in L2/3. For somatic GCaMP recordings, the FOV was either 480 × 480 µm or 320 x 320 µm, recorded at ∼10 Hz (2 planes, 40 µm apart). Laser power measured after the objective was between 18-37 and 44-52 mW at 920 nm and 1070 nm, respectively. For each mouse, the objective optical axis was aligned perpendicularly to the imaging window.

Ambient noise in the sound-insulated two-photon recording cage was 39.0 ± 0.02 dB SPL and 39.2 ± 0.06 dB SPL during the axonal recordings, with SPL values calculated within the mice hearing frequency range (2-80 kHz). By comparison, broadband white noise stimuli (2-80 kHz) reached the microphone placed at the location of the mice head at 48.0 ± 0.02 dB SPL, as expected from calculations based on sound attenuation in the air. Bandlimited white noise (2-20 kHz) reached the center of the perimetric device at ∼66 dB SPL. Scanning noise generates noise at 8 kHz. At this frequency specifically, ambient noise was 21.6 ± 0.09 dB SPL, scanning noise was 26.9 ± 0.46 dB SPL and broadband white noise stimuli was 30.62 ± 0.31 dB SPL. Values are mean ± s.d., estimated from 3 measurements. Scanning a 320 µm^2^ FOV (used for somata imaging during multimodal experiments) generated noise at 42.5 dB SPL and 39.6 dB SPL specifically at 8 kHz. Bandlimited white noise used in these experiments were estimated to reach the mouse head at ∼66 and 48 dB SPL (generated at 85 and 65 dB SPL, calibrated with a microphone 2.5 cm away from the speaker).

### Speaker and LED array

The speaker and LED array was built in-house. Its design is available through the Champalimaud Hardware Platform (http://www.cf-hw.org). Thirty-nine speakers (MULTICOMP PRO, # ABS-239-RC) and LEDs were mounted on a 3D-printed resin framework. Each position containing both a speaker and a LED were 10 degrees apart in azimuth and 20 degrees apart in elevation, thereby taking the shape of a spherical section spanning 120 degrees in azimuth and 40 degrees in elevation, with a radius of 18 cm.

We quantified reverberation in the speaker array. We applied a white noise pulse (20 ms, 2-80 kHz) and recorded the sound from a microphone placed in the center of the device. The power of the sound after offset decayed to half of the power in 3.7 ± 0.7 ms (exponential fit, mean ± s.d. across speakers, measure repeated three times). We fitted the decay of the log power of the sound after sound offset over time with a linear function and quantified the rate of this decay by the time taken for the reverberating sound to decay 20 dB (20-dB reverberating time, RT20). RT20 was 19.4 ± 0.74 ms (mean ± s.d. across speakers, measure repeated three times). Similar values were obtained for the RT20 of the 2-20 kHz frequency band.

#### Auditory stimulation

Auditory stimuli were equalized white noise (2-20 or 2-80 kHz). Each speaker was individually calibrated to flatten its frequency response using a Brüel & Kjær Free-field 1/4-inch microphone placed 2.5 cm from the speaker. All speakers were regularly calibrated. Standard deviation of the power across frequencies for each speaker typically ranged from 0.97-1.57 dB (mean across speakers, 1.19 dB).

Sounds were digitized and amplified using a custom-built sound card and amplifier (192 kHz sampling rate, 24 bits precision; https://www.cf-hw.org/harp/harp-devices#h.p_lrxK9t3VI9s0). All the speaker-specific waveforms obtained from the calibration were stored in different channels of the sound card to minimize delay for sound stimulus presentation and applied for each speaker on a trial-by-trial basis.

The speakers playing the sound were switched 100 ms before sound presentation using three clickless audio switches (MAXIM, MAX4910; HARP audio switch https://www.cf-hw.org/harp/audio-switch), controlling 12 to 15 speakers each.

For spatial tuning in AC axons, sound stimulation consisted of five 200 ms bursts with 10 ms raised-cosine onset/offset ramps (1 s total). Unless otherwise mentioned, band-limited white noise (2-20 kHz) were played at 85 dB SPL and reached the mouse head at ∼66 dB SPL. Broad-band white noise (2-80 kHz) were played at 65 dB SPL and reached the mouse head at ∼48 dB SPL (loudness was calibrated with a microphone at 2.5 cm away from the speakers).

For audio-visual experiments, sound stimulation consisted of five 100 ms-long bursts (with 10 ms raised-cosine onset/offset ramps) with a 50% duty cycle (1s in total).

For the experiments in Figure 7 a single loudspeaker (MULTICOMP PRO, # ABS-239-RC) was mounted on motorized rotation stage controlled by a stepper motor (8MR151 and 8SMC5-USB, respectively standa.it) to present sound from –90 (left, ipsi-) to +100° (right, contra-lateral) around the frontal position, at a radius of 18 cm.

#### Visual stimulation

The RGB LEDs were controlled using an LED controller board built in-house (HARP RGB LED; https://www.cf-hw.org/harp/led-arrays). Light diffusers (5 mm-thick acrylic fitted in a 3D-printed frame, not represented in Figure 1a) were positioned in front of each LEDs. In the absence of visual stimulation, all the LEDs were set at a dim blue light level (0.25 cd/m^2^) to provide some background light. For V1 neurons’ RF mapping, LEDs were flashing white light at 3 Hz (50% duty cycle) at a luminosity of 11.4 cd/m^2^. For audio-visual experiments using the perimetric device, LEDs were flashing white light in synchrony with the auditory stimulation at 5 Hz for 1 s (50% duty cycle) at luminosities of 2, 11.4 or 20.8 cd/m^2^.

HARP Audio switches, HARP RGB LED board and HARP sound card were synchronized using a HARP clock synchronizer (https://www.cf-hw.org/harp/clock-sync). We ensured auditory and visual stimuli were synchronized using an oscilloscope to verify the delays.

Stimulus sequences were generated using MATLAB (The Mathworks, Natick, MA). All the components in the 3D array were controlled using Bonsai ^76^. Auditory and visual stimulation positions were pseudo-randomized and independent between speaker and LED.

### Histology

After completion of imaging experiments, mice were deeply anesthetized, transcardially perfused, and fixed overnight with 4% paraformaldehyde in 0.1 M phosphate buffer, pH 7.4. The brain was cut into 50-µm-thick coronal sections with a vibrating slicer (Leica). GCaMP was amplified using a polyclonal anti-GFP antibody (ThermoFisher, catalog #A-6455) and an Alexa Fluor 488-conjugated secondary antibody (ThermoFisher, catalog #A-11008) and counterstained with DAPI. Images were taken using a slide scanner (Zeiss AxioImager M2) connected to a sCMOS camera (Hamamatsu ORCA-Flash 4.0) with either a 10× (0.45 NA) or 20× (0.80 NA) objective (Zeiss). Retrogradely labeled cell bodies from AC in V1 were absent. We used QuickNII and VisuAlign ^77^ to align the brain sections to the Allen Mouse Brain Atlas and identify the boundaries of cortical areas.

### Videography

We recorded videos of the mouse eye/face during two-photon imaging. We used a CMOS camera (Flea3 USB3 Vision, PointGrey) mounted with a telephoto lens (Navitar Zoom 7000) and a high-pass filter. The eye was imaged using scattered 920nm light during two-photon imaging. For full-face movies, a high-power infrared LED (850 nm, Roithner LaserTechnick) was used to illuminate the face. Videos were recorded at ∼20 Hz and the acquisition was synchronized with two-photon recording using a common TTL trigger sent by the LED controller board.

Pupil was delineated post-hoc using DeepLabCut ^78^. In ∼200 video frames, eight points along the edge of the pupil were manually annotated (every 45°). Then a resnet-based convolutional neural network was trained to predict the location of these markers. The trained network was then manually evaluated and retrained by manually correcting labels from a subset of poorly predicted frames. The final network was then used to track the eight labels on every video frame. In video frames with average label placement probability > 0.9, with at least 6 markers with probability > 0.8, and where basic assumption were not violated – for instance the top eye label should have higher y coordinates than the bottom one – an ellipse was fitted to trustworthy the labels and pupil area was calculated.

Principal components of motion energy from full-face videos were extracted using the singular value decomposition provided by the Facemap algorithm ^79^ (https://github.com/MouseLand/facemap).

#### Two-photon data analysis

##### Preprocessing of axonal data

Motion correction and regions of interest (ROI) segmentation was performed using Suite2p ^80^. We used the ‘anatomical’ parameter, such that suite2p placed ROIs over varicosities (likely presynaptic axonal boutons, referred as boutons for simplicity) rather than axon segments. ROIs were manually added over boutons that were not picked up by Suite2p. Under the ‘anatomical’ detection, detected ROIs always encompassed single boutons. In a subset of experiments (5/8 AC mice, 2-20 kHz white noise), ROIs were detected using the ‘functional’ parameter in suite2p, yielding detection of ROIs containing several boutons. We split responsive ROIs into (multiple) boutons using a watershed procedure and all the resulting single bouton ROI were assigned the same *ΔF/F* trace.

We calculated *ΔF/F = (F-F_0_)/F_0_*, where F is the average fluorescence of all pixels in each ROI and F_0_ is the fluorescence of the ROI during the trial baseline.

Axonal ROIs were considered responsive if the strongest median response across all positions was different from the baseline (two-sided Wilcoxon signed-rank test for paired sample test, α = 0.01) and the response amplitude was larger than 0.15 ΔF/F_0_. Results where qualitatively identical when selecting responsive ROIs with a bootstrapping method (strongest median response superior to 95% confidence interval of the baseline, and amplitude of response larger than 0.15 ΔF/F_0_).

The fraction of responsive boutons was averaged across sessions per mice. In Figure 2d, all responsive boutons from all mice were used.

##### Comparison of Azimuth and Elevation modulation

To compare how responses axons depended on azimuth and elevation of the speaker we downsampled azimuth positions to simulate a 40°x 40° isotropic speaker array. Azimuth positions were downsampled to all possible combinations of three positions covering 40° and 20° steps. The number of space-sensitive boutons was determined using a two-way ANOVA for each of and averaged across the 9 possible virtual arrays. Data was further averaged across positions of the same mouse and significant difference in the fractions of azimuth, elevation and interaction-sensitive boutons was tested using a one-way ANOVA.

##### Spatial Modulation Index (SMI)

SMI quantifies the relative proportion of the energy of the RF that could not be explained by the spatial average response and was calculated as previously described ^81^

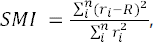

where r_i_ is the average response to the stimulus in position i, n is the total number of positions and R is the average response to all stimuli.

##### Decoding sound and light stimuli location from axonal recordings

To decode the location of the stimulus from the recorded axonal activity we used a naive Bayesian decoder. Because a Bayesian decoding assumes independence of the data, we first grouped together the ROIs that belong to the same axon. We used a correlation-based method to distinguish ROIs that were part of the same axon and those that were not ^46^. We first identified boutons that had visible axon shaft between them to calculate correlation values for boutons from the same axon. We determined that 58% of the boutons from the same axon had a Pearson’s correlation value equal to or larger than 0.3 while less than 0.5% of the general population did. Based on this correlation value we selected all the pairs that were from the same axon and seeded a cluster with one randomly selected pair. The next randomly selected pair could be correlated with a coefficient over this value with only one of the ROIs with the existing cluster, in which case it joined the cluster, or not, in which case it seeded a second cluster. We iterated this procedure until all pairs were assigned. Activity in each cluster was assigned to the ROI with the largest mean ΔF/F. In sessions where this method yielded more than 10 axons, we applied a naïve Bayesian decoder approach.

For each selected session, we trained the decoder as follows. The probability that the stimulus at a given trial was coming from the location s is given by Bayes’ theorem:

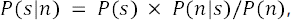

where n is the response of the entire axonal population. Assuming statistical independence of activity in the N axon recorded, we derived the likelihood function *P*(n/s) as:

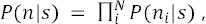

where n_i_ is the response of the axon i.

Responses were modeled using a normal distribution and thus, for all axon i and stimulus s:

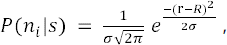

where, for axon *i*, *σ* is the standard deviation of response to the stimulus s, r is the axon’s response to the stimulus and R is the mean response to the stimulus s.

Therefore, the log-likelihood function is defined as:

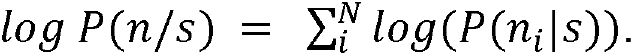

The decoded position is the maximum likelihood estimate ŝ i.e., the position that maximizes the log likelihood.

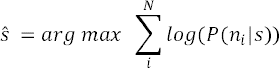

For each trial, likelihood was computed for all 39 possible positions and the maximum was used to decode the stimulus location. In Figure 3b and Supplementary Figure 3a, decoding in azimuth was considered correct if it fell in the actual stimulus position ± 10° in azimuth, regardless of elevation.

The likelihood distribution and the mean and standard deviation of the responses were computed on different sets of trials using a 5-fold cross-validation procedure. Decoding accuracies were averaged over the 5 runs to extract the decoding performance per stimulus position for one two-photon recording session. Accuracies were then averaged across stimulus location and recording sessions from the same mouse. Decoding accuracies for the shuffle data was obtained using the same procedure, but for data where axon identity was permuted, thereby preserving the total magnitude of the response per stimulus location.

For calculating the decoding performance relative to the V1 neuron’s population RF azimuth center (Figure 4c,d), the decoding accuracy was measured relative to the absolute difference between the azimuth location of the stimulus and V1’s population RF azimuth center to the stimulus location. Data for a given distance between speaker location and V1 neuron’s population RF were kept if generated in 5 or more sessions.

For calculating the distance between the decoded and actual stimulus position vs. number of axons used (Figure 3c), we pooled axons from all the recording sessions and randomly drew a given number of axons without replacement (not to violate the assumption of data independence). The distance between the actual and decoded stimulus position was calculated on a single trial basis and averaged across trial types and runs. This procedure with random sampling was repeated 100 times to evaluate the mean and 95% confidence interval of the distance between decoded and actual stimulus position for a given number of axons.

To calculate the accuracy as a function of V1 position from which the axons were recorded from (Figure 5b), we randomly drew the n_min_ – the minimal number of axons across sessions – 10 times per session and averaged the decoding accuracy across resampling and calculated the moving average over 20° in azimuth in 5° steps.

##### Peak azimuth

We selected reliable boutons to measure their spatial profile. Trials of each trial type were randomly split in two. Boutons were considered reliable if the Pearson’s pairwise correlation coefficient between the 2 halves was > 0.3. The preferred stimulus azimuth position was estimated from the responses averaged across all trials and elevations. The session’s peak azimuth was extracted by taking the mean of the distribution of the bouton’s preferred azimuthal position. This procedure was repeated 100 times to estimate the median and 95% confidence interval of the peak azimuth per session. Only sessions with at least 10 reliable boutons were included.

##### Spatial tuning curves

Trials of each trial type were randomly split in two and boutons’ spatial tuning was cross-validated if Pearson’s pairwise correlation coefficient between the 2 halves was > 0.3. For each cross-validated bouton, the preferred stimulus position (maximum mean response across trials) was obtained from the first random half of the trials and responses to the second random half were normalized to that maximum. Preferred azimuth was obtained from by averaging responses across elevation, and vice versa for elevation. Boutons with the same preferred azimuth/elevation were pooled and normalized responses of the second random were plotted. This procedure was repeated 100 times to estimate the mean and 95% confidence interval of the tuning curves. Responsive boutons across all recording sessions from all mice were used.

##### Measurement of V1 L2/3 somata population receptive fields

Full field-of-view responses were baseline subtracted (1 s to frame before LED stimulus onset) and averaged across repetitions. Responses during the stimulus window were time-averaged (1st frame after stimulus onset to 1st frame after stimulus offset, 1 s) to obtain a map of stimulus response. The stimulus response map R(az, el), where az and el are the coordinates azimuth and elevation, respectively, was fitted with a two-dimensional Gaussian. In audiovisual experiments (Figure 6), we then placed the visual stimuli in the center of the population RF.

##### Preprocessing of somatic recordings

For GCaMP6f somatic recordings (Figure 6), motion correction and regions of interest (ROI) segmentation was performed using Suite2p like for axonal data ^80^. Data was manually curated. Fluorescence within each ROI was neuropil corrected using (*F* = *F* – 0.7, x *Fenu,* where Fneu is the neuropil’s fluorescence).

For the analysis of the magnitude of the audiovisual modulation, we excluded neurons with high variability in their response (s.d of the response > 10, <1 % of all neurons) All somas were included for the decoder analysis.

##### Decoding sound location from somatic V1 recordings

A naïve Bayesian decoder was built from somatic recordings data in each recording session. Speaker position in individual AV trials were decoded using the maximum likelihood estimate as for axonal data (see above). We used a leave-one-out cross-validation and averaged the decoded distance to actual speaker position in the AV trial over the 7 runs, and across sessions and mice. Shuffle data was obtained by randomizing the trial labels without replacement 100 times.

