## Supplemental Figures for "Auditory cortex conveys non-topographic sound localization signals to visual cortex"

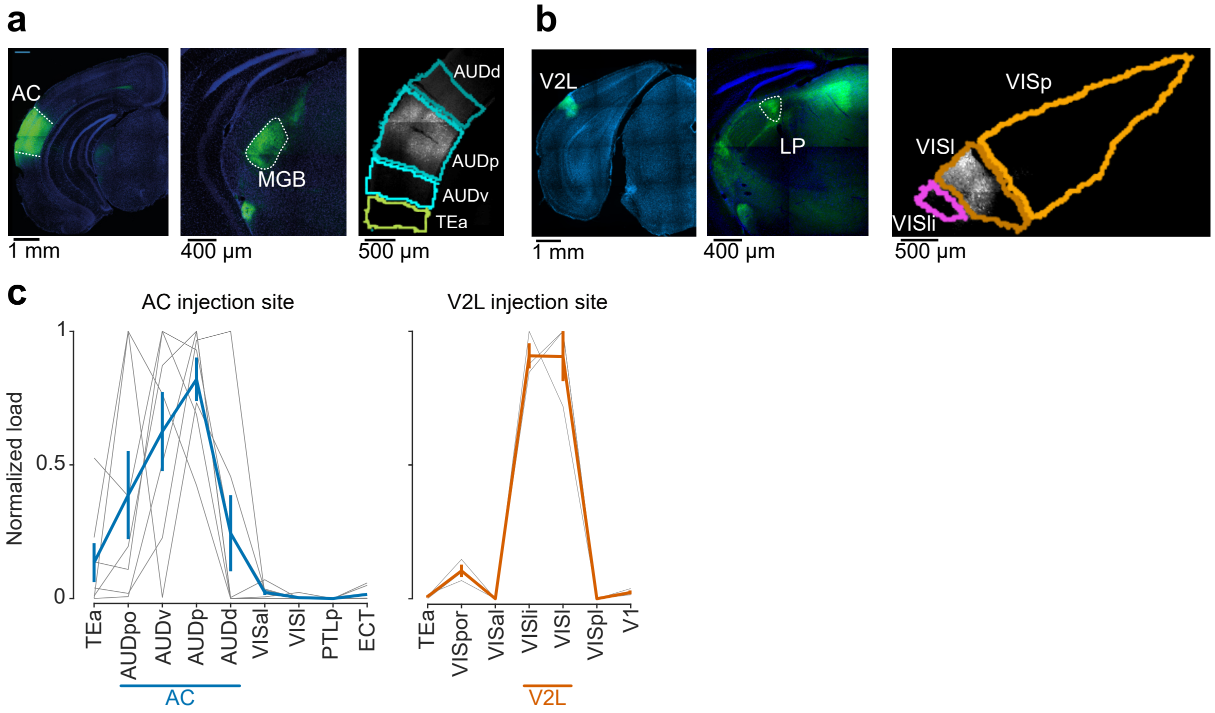


**Supplementary Figure 1. Histological analysis of AC and V2L -injected mice. a** Example coronal sections of an AC-injected brain. Left, Section through the injection site in the AC cortex. Center, axon projections in the thalamus . Right, sections were registered to the Allen mouse brain atlas to estimate the relative GCaMP expression across the AC and neighboring regions. **b** Same for an example V2L-injected brain. **c** Quantification of fluorescence in different areas around the injection site for AC- (blue, left) and V2L-injected mice (yellow, right). Gray lines, individual mice; colored line, average ± s.e.m. across mice. n = 7 AC-injected mice and n = 3 V2L-injected mice. Auditory areas: AC, auditory cortex; AUDpo, posterior auditory cortex; AUDd, dorsal auditory cortex; AUDp, primary auditory cortex; AUDv, ventral auditory cortex; MGB, medial geniculate body; Visual areas: V2L, higher-order lateral visual area; V1, primary visual cortex; VISpor, postrhinal area; VISal, anterolateral visual area; VISl, lateral visual area; VISl, lareomedial visual area; VISli, laterointermediate visual area; VISpl, posterolateral visual area; LP, lateroposterior nucleaus; Others: TEa, temporal association areas; PTLp, posterior parietal association area; ECT, ectorhinal area.

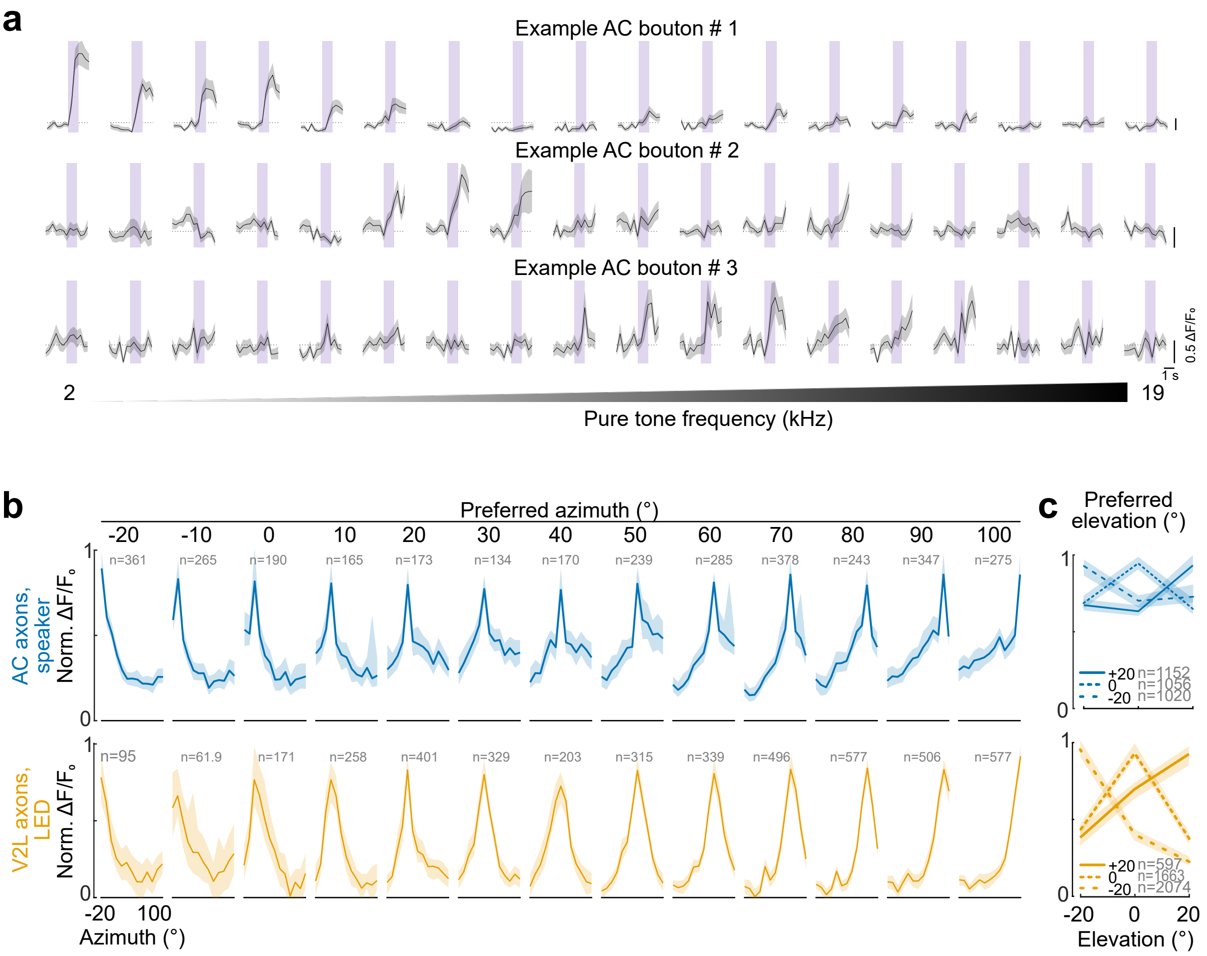


**Supplementary Figure 2. Frequency and spatial tuning curves in AC→V1 inputs. a** Average responses to different pure tones (2-19 kHz, 1 kHz step, purple shaded area) from three example AC boutons recorded in the same imaging session. **b** Normalized azimuth tuning curves. Each plot is the mean of all the cross-validated boutons preferring sounds located at a given azimuthal position (see Methods). Colored lines, mean across boutons; shaded area, 95% confidence interval. The mean number of boutons is indicated in gray. **c** Same as **b** for elevation.

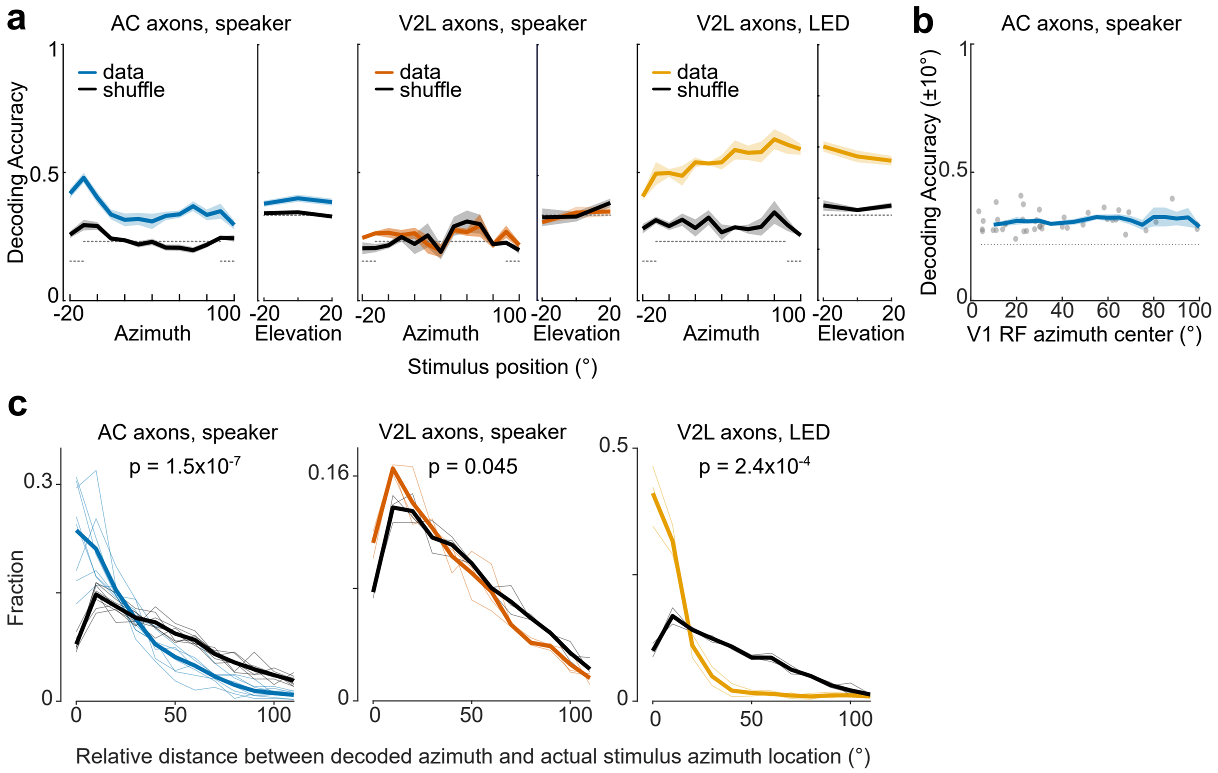


**Supplementary Figure 3. Decoding of stimulus location from AC and V2L inputs in V1.** **a** Decoding accuracy as a function of stimulus azimuthal (left insets) and elevation location (right insets) in auditory responses in AC axons (left) and V2L axons (middle) and in visual responses in V2L axons (right). In azimuth, decoding accuracy is considered accurate if the estimated location is within ± 10° of the actual location. Colored lines, average decoding accuracy across mice; black lines, decoding accuracy using shuffled data. Shaded area is the s.e.m across mice. n = 8 AC- and n = 3 V2L-injected mice. Dotted lines are chance level. **b** Overall decoding accuracy as a function of imaging position in V1, with matched number of axons per position. V1 RF azimuth center measured using LED responses in jRGECO1a+ somatas in L2/3 of V1. Dots, individual sessions; curve and shaded area, moving average and s.e.m performance calculated over 10° bins. One-Way ANOVA: $F(19,87)=0.55$, $p=0.9$. **c** Distribution of the decoded position relative to actual stimulus position. Thin lines are individual mice, thick lines are mice average per group. Two-way repeated measures ANOVA, p values for interactions are reported.

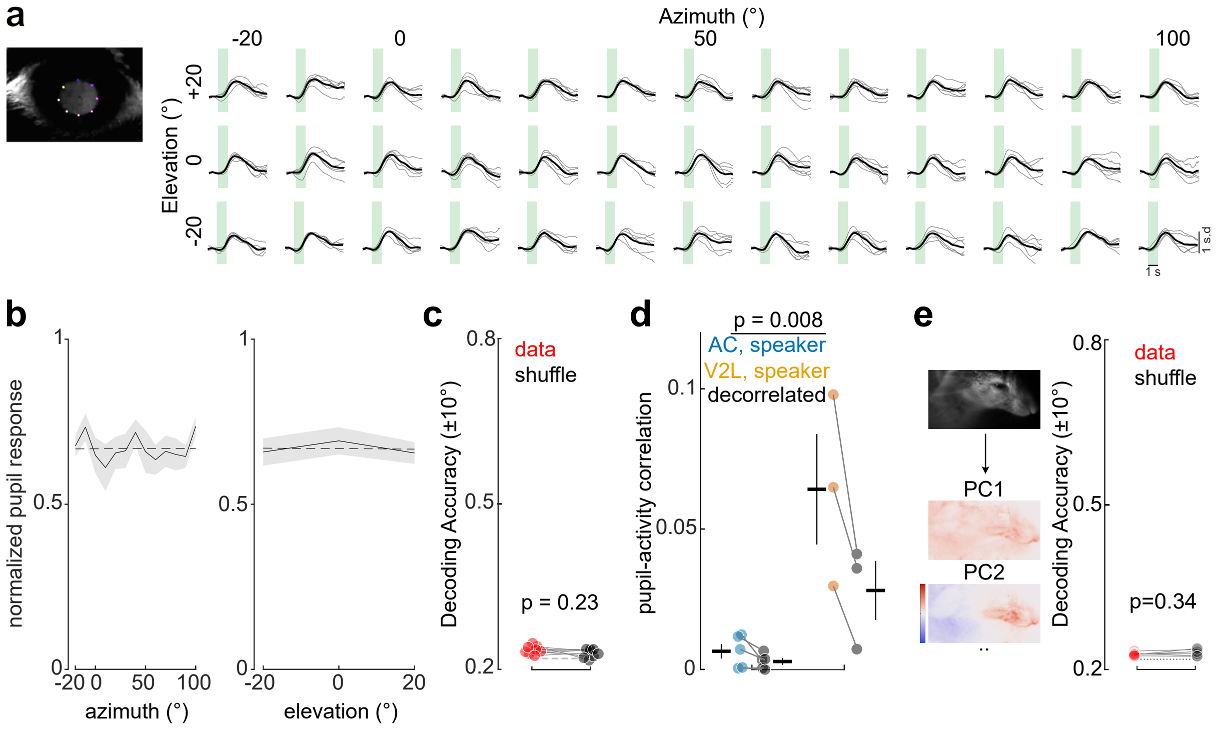


**Suplementary Figure 4. Sound-induced behavioral responses are similar across space.** **a** Left, Automated pupil measurements. Example video frame from one session plotted on the right. Colored dots, labels generated by Deeplabcut. Right, Sound-induced pupil responses for sounds played at different positions of the speaker array for an example mouse. Green shaded area, auditory white noise; Gray lines, sessions; Black lines, average across sessions of an example mouse. **b** Normalized pupil responses across azimuthal (left) and elevation (right) showed no systemic bias (One-way ANOVA: p = 0.82 and p = 0.76 for azimuth and elevation, respectively, n = 8 mice: 5 AC- and 3 V2L-injected mice). Data is mean across mice ± s.e.m.; dotted line, linear fit. **c** Accuracy of a naive Bayesian decoder using pupil response data (red) versus the trial shuffle data (black). Accuracy averaged across sessions. Circles, mice; dotted line, chance. Two-sided paired t-test: n = 8 mice. **d** Pearson's correlation between pupil and auditory responses in AC (blue) and V2L boutons (yellow). Dots, average correlations across boutons per mice (orange). Crosses, mean ± s.e.m; gray/black, corresponding trial-shuffle data ('decorrelated'). In AC-injected mice, pupil-bouton pairwise correlations in auditory responses were not significantly different between actual (blue) and trial-shuffle data (black; two-sided paired t-test: p = 0.09, n = 8 mice). Correlations of auditory responses with pupil size were larger in V2L- than in AC-injected mice (two-sided t-test). **e** Left, top, average of all frame of a movie from an example movie (CBA mouse). Middle and bottom, spatial masks of the top two principal components (PC) of the motion energy movie. Color scale indicates low-motion energy (blue) to high-motion energy (red). PC1 captures motion anywhere on the mouse's face while PC2 highlights movement in the snout area. Right, decoding accuracy of sound location from the 500 first PCs of the face movie (red) was not significantly higher than that of the trial shuffle data (black; two-sided paired t-test: p = 0.34, n = 6 CBA mice stimulated with 2-80 kHz sounds). Dots, accuracy per mouse, averaged across session; dotted line, chance level.

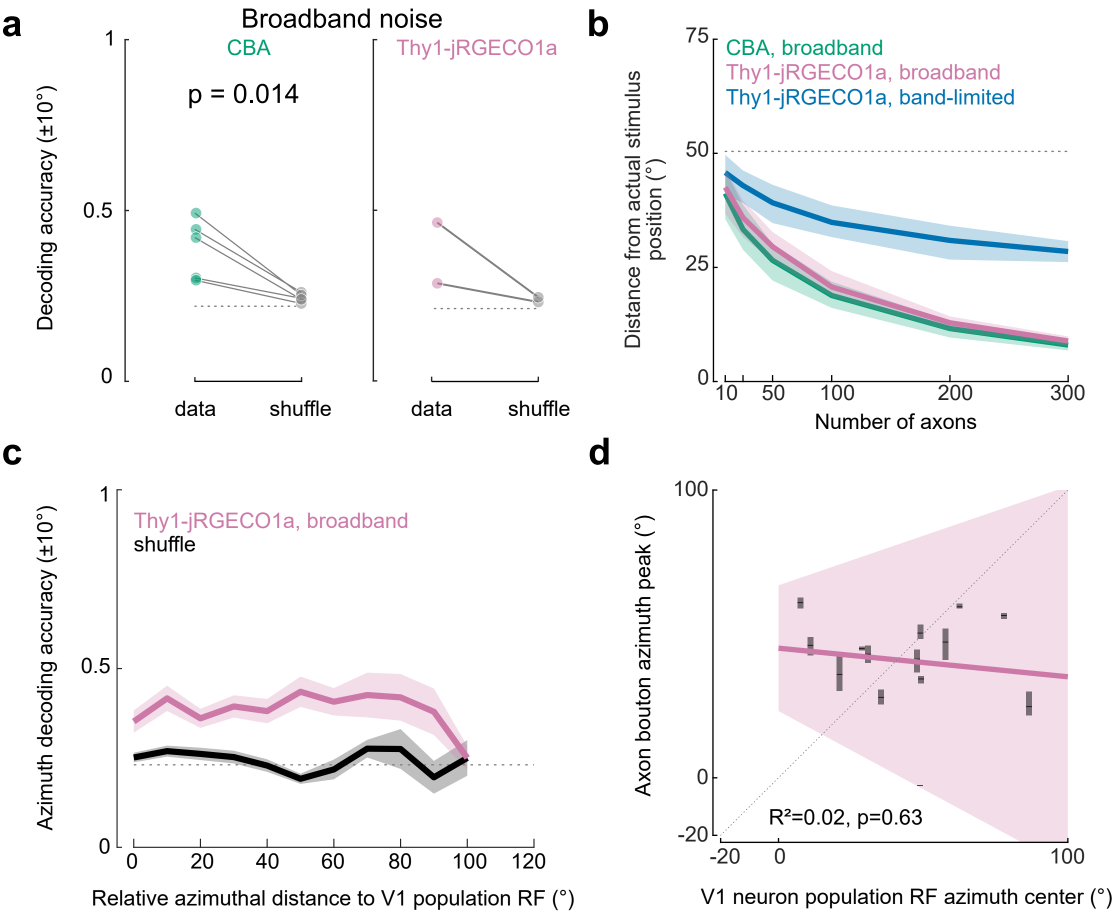


**Supplementary Figure 5. Including high-frequencies improves decoding of sound location from AC→V1 inputs but did not reveal a topographic organization. a,** Azimuth stimulus location decoding accuracy using high-frequency containing sound in CBA (left) and Thy1-jRGECO1a mice (right). Colored circles, average across sessions for each individual mouse; gray circles, shuffled data from individual sessions, averaged across session per mouse. Dotted lines denote chance. Two-sided paired t-test: n = 5 CBA mice, 30 sessions and n = 2 Thy1-jRGECO1a mice, 14 sessions. **b** Decoding accuracy as a function of number of axons for Thy1-jRGECO1a mice, band-limited noise (blue), broadband noise (purple) and CBA mice, broadband noise (green-blue). Lines are median across resampling interations; shaded areas are 95% confidence intervals; dotted line, chance. **c** Decoding performance as a function of the distance between stimulus location and V1 population RF center in Thy1-jRGECO1a mice using broadband sounds. Data is mean ± s.e.m.; dotted line, chance. One-way ANOVA: $F(12,10) = 0.08$, $p = 0.71$. **d** Mean peak azimuth of the sound evoked response across AC boutons as a function of the population RF center of V1 neurons for each imaging session. Ticks, median; grey shading, 95% confidence interval. Colored lines, linear fits of the median values; colored shading, 95% confidence interval; dashed line, identity line. n = 14 sessions, 2 mice.

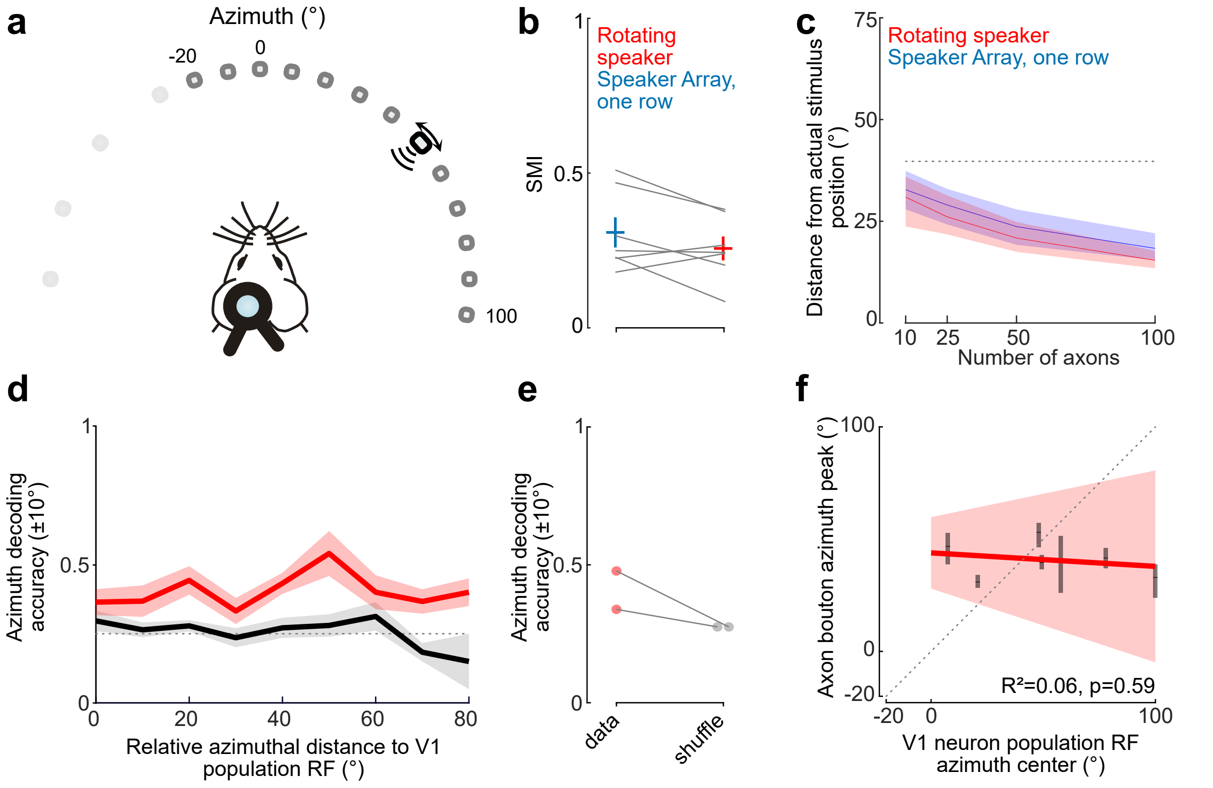


**Supplementary Figure 6. Sound location specific signals in AC inputs to V1 measured using a single loudspeaker. a** A single loudspeaker (black rectangle, 2-80 kHz white noise) was moved between -20° to +100° in azimuth, 10° spacing (dark gray positions; 0° elevation). Light gray positions represent positions sampled during the same experiment, analyzed separately (see **Figure 5**). **b** Similar SMI values were obtained with a rotating speaker and with the middle row (0° elevation) of the speaker and LED array (two-sided paired t-test: p = 0.15, n = 7 imaging sessions from 2 mice). **c** Speaker location decoding accuracy as a function of number of axons used was similar across both conditions. Lines and shaded area are means and 95% confidence intervals. Gray dotted line indicate chance level. **d** Decoding performance using the rotating speaker does not depend on V1 retinotopy (One-way ANOVA: F(9,46)=1.12, p = 0.37, n = 7 sessions from 2 mice). Red: observed data; black: shuffle data. Data is mean ± s.e.m across sessions. **e** Average azimuth decoding accuracy for the 2 mice (average across sessions). Dots, mice; lines indicate paired data. **f** AC bouton mean peak azimuth are not topographically organized according to V1 retinotopy (n = 7 sessions from 2 mice).
